# Structure of the human inner kinetochore bound to a centromeric CENP-A nucleosome

**DOI:** 10.1101/2022.01.07.475394

**Authors:** Stanislau Yatskevich, Kyle W. Muir, Dom Bellini, Ziguo Zhang, Jing Yang, Thomas Tischer, Masa Predin, Tom Dendooven, Stephen H. McLaughlin, David Barford

**Author notes:** These authors contributed equally to this work (Stanislau Yatskevich, Kyle W. Muir).

## Abstract

Accurate chromosome segregation, controlled by kinetochore-mediated chromatid attachments to the mitotic spindle, ensures the faithful inheritance of genetic information. Kinetochores assemble onto specialized CENP-A nucleosomes (CENP-A^Nuc^) of centromeric chromatin. In humans, this is mostly organized as thousands of copies of an ∼171 bp *α*-satellite repeat. Here, we describe the cryo-EM structure of the human inner kinetochore CCAN (Constitutive Centromere Associated Network) complex bound to CENP-A^Nuc^ reconstituted onto α-satellite DNA. CCAN forms edge-on contacts with CENP-A^Nuc^, while a linker DNA segment of the α-satellite repeat emerges from the fully-wrapped end of the nucleosome to thread through the central CENP-LN channel which tightly grips the DNA. The CENP-TWSX histone-fold module, together with CENP-HIK^Head^, further augments DNA binding and partially wraps the linker DNA in a manner reminiscent of canonical nucleosomes. Our study suggests that the topological entrapment of the *α*-satellite repeat linker DNA by CCAN provides a robust mechanism by which the kinetochore withstands the pushing and pulling of centromeres associated with chromosome congression and segregation forces.

**One-Sentence Summary:** The human inner kinetochore CCAN complex tightly grips the linker DNA of the α-satellite CENP-A nucleosome.

## Main Text

The centromere is a specialized genetic locus that interacts with the mitotic spindle to facilitate chromosome segregation. Centromeric loci of humans and other primates comprise multiple copies of the conserved ∼171 bp *α*-satellite repeat that contains the histone H3 variant CENP-A (*1, 2*). Kinetochores assemble specifically onto centromeres to form large macromolecular machines that directly mediate chromosome segregation (*3–5*). A critical function of the kinetochore is to generate a load-bearing attachment between chromosomes and the spindle apparatus. How this function is achieved at the molecular level is unclear.

Throughout the cell cycle, CENP-A nucleosomes specify the constitutive association of the 16-subunit inner kinetochore CCAN complex with centromeric chromatin, whereas the ten-subunit KMN network of the outer kinetochore assembles onto CCAN during mitosis to couple CCAN to microtubules for spindle-mediated chromosome movement. Two human CCAN proteins, CENP-C and CENP-N, have been observed to directly interact with CENP-A^Nuc^ (*6, 7*). CENP-C interacts with the C-terminus of CENP-A, in conjunction with histones H2A and H2B, whereas structures of the N-terminal domain of human and chicken CENP-N in complex with CENP-A^Nuc^ revealed a CENP-N-binding site on the L1 loop and the adjacent DNA gyre of CENP-A^Nuc^ (*8–12*). The 17 bp B-box motif that interacts with CENP-B is the only DNA sequence within the *α*-satellite repeat that is specifically recognized by a centromere-associated protein, however B-box motifs are absent from the Y-chromosome, and the genomes of new world monkeys. CENP-B, although non-essential (*13–15*), increases the fidelity of chromosome segregation (*16*).

In budding yeast, a single kinetochore complex associates with a ‘point’ centromere accommodating a sole CENP-A nucleosome. In contrast, the regional kinetochores of humans and other metazoans are generated from multiple copies of the CCAN and KMN network complexes to form large disk-like structures. Here, we determined the cryo-EM reconstruction of a CCAN-CENP-A^Nuc^ complex using a native 171 bp *α*-satellite repeat DNA. This represents the repeating modular unit of a human inner kinetochore. In our structure, the emerging extranucleosomal DNA, that links adjacent nucleosomes, threads through a DNA-binding tunnel in the CCAN molecule. The topological capture of centromeric DNA by human kinetochores provides an explanation for how load-bearing attachments between centromeres and spindle microtubules can be formed (*11, 12, 17, 18*).

## Results

### CCAN is assembled from a network of interdependent modules

Human CCAN comprises four defined modules: CENP-LN, CENP-HIKM, CENP-OPQUR and CENP-TWSX. These modules are additionally linked by the largely disordered CENP-C that interacts directly with CENP-LN and CENP-HIKM (*8, 18, 19*). We expressed and purified all CCAN sub-complexes recombinantly, using the N-terminal half of CENP-C (CENP-C^N^) that contains all known CCAN binding sites as well as the central domain essential for centromere localization (*20, 21*) (**fig. S1A**). Initially, we focused on the 11-subunit CCAN core complex (CENP-OPQUR-LN-HIKM) which excluded the CENP-TWSX module, and could assemble in the absence of CENP-C^N^ (termed CCAN^ΔCT^) (**fig. S1B**). We determined the cryo-EM structure of CCAN^ΔCT^ at 3.2 Å resolution (**Fig. 1, fig. S2 and Table S1**). The cross-linked structure was identical to a non-cross-linked reconstruction obtained at a lower resolution of 8 Å that included CENP-C^N^ (termed CCAN^ΔT^) (**fig. S2D**). We also determined crystal structures of the CENP-OPQUR and CENP-HIK^Head^ modules (**fig. S2E and Table S2**).

**Figure 1.**
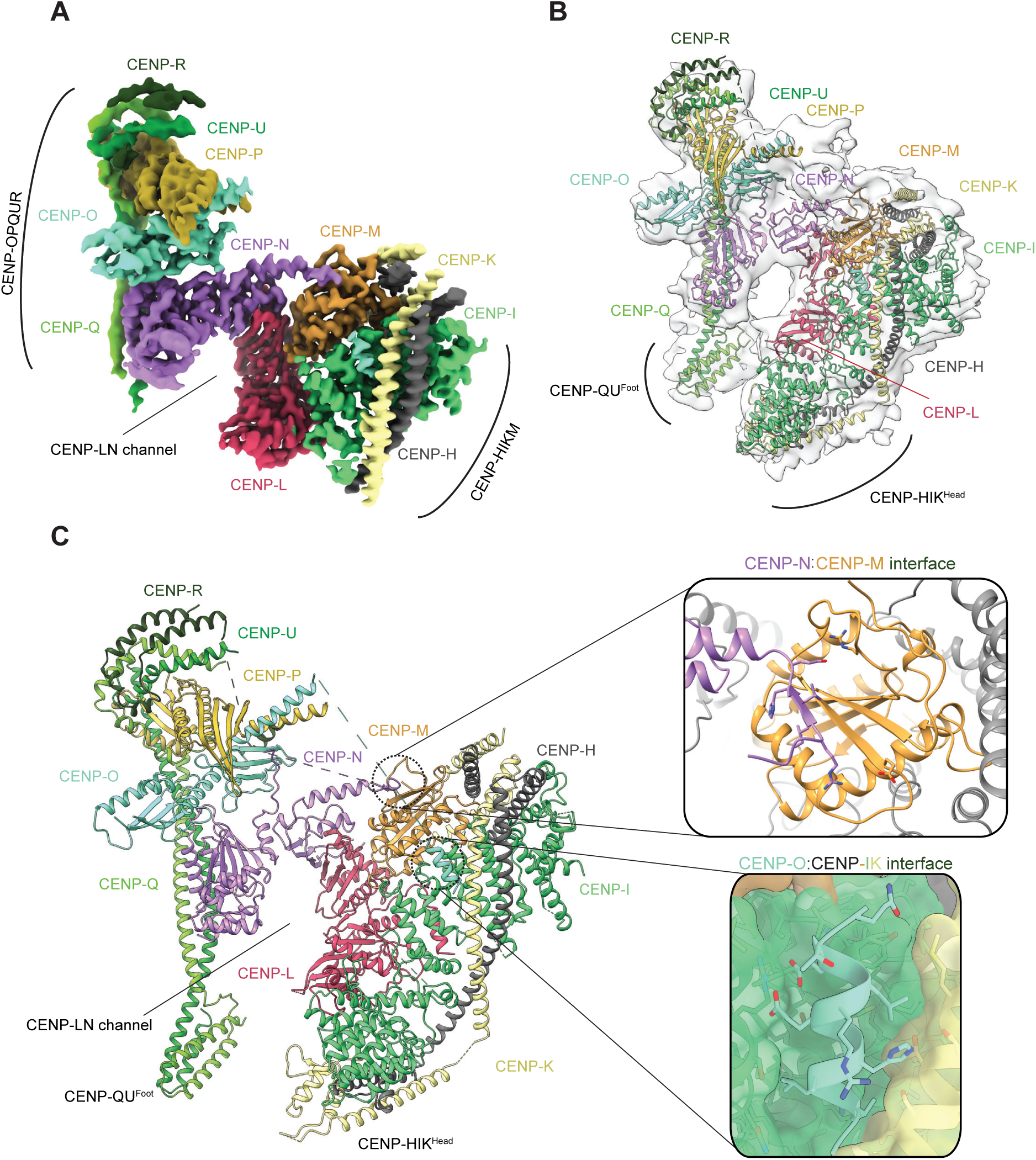
CCAN is assembled from a network of interdependent modules. (**A, B**) Cryo-EM density map (**A**) and molecular architecture (**B**) of the apo-CCAN^ΔCT^ fitted into the cryo-EM map with resolved OPQUR^Foot^ and HIK^Head^. (**C**) Complete atomic model of the apo-CCAN^ΔCT^ with details of the CENP-N-CENP-M interface as well as CENP-O binding to the CENP-HI pocket are shown as insets.

The architecture of CCAN^ΔCT^ resembles budding yeast CCAN (*Sc*CCAN) (*22, 23*) with CENP-OPQUR and CENP-HIKM ‘lobes’ assembled on either flank of a central CENP-LN module (**Fig. 1**). The arc-like CENP-LN module is the structural keystone of CCAN^ΔCT^, and generates a deep channel that is extended on one side by the N-terminal coiled-coil of CENP-QU. The RWD domains of CENP-O and CENP-P, with the C-terminal regions of CENP-QU and CENP-R, organize into a ‘cap’ domain above CENP-N. In human CCAN^ΔCT^, CENP-R partially substitutes for the budding yeast Nkp1 and Nkp2 subunits. CENP-M, that is also unique to vertebrate CCAN, adopts a pseudo-GTPase-fold and forms a stable complex with CENP-HIK (*24*). CENP-M makes extensive contacts with the HEAT domain of CENP-I and the N-terminal helical bundle of CENP-H and CENP-K, and in the context of CCAN, with CENP-L (**Fig. 1**). Its position, wedged into a gap separating the C-terminal domains of CENP-I and CENP-L, causes CENP-L, with CENP-HIK, to rotate about the CENP-N-CENP-L interface towards CENP-OPQUR, thereby narrowing the CENP-LN channel compared to *Sc*CCAN (*23*). CENP-M, when compared to other GTPases, has a small deletion that removes the switch I region and an adjacent *β*-strand (*24*). A conserved *β*-strand of CENP-N occupies this position to augment the central *β*-sheet of CENP-M (**Fig. 1C**). An N-terminal *α*-helix of CENP-O, connected through a flexible linker to CENP-OPQUR, interacts with a conserved pocket formed by CENP-I and CENP-K (**Fig. 1C**). Disrupting this interaction by deleting the N-terminal 35 residues of CENP-O destabilizes CCAN^ΔCT^ assembly, as does mutating the five inter-domain residues of CENP-N that bind CENP-M (**fig. S3A, B**).

Two peripheral and conformationally flexible regions of CCAN are poorly resolved in the cryo-EM reconstruction: the CENP-HIK^Head^ and N-terminal region of CENP-QU (CENP-QU^Foot^) (**Fig. 1**). Focused 3D classification of CENP-HIK^Head^ identified multiple poses, two of which readily accommodate the CENP-HIK^Head^ crystal structure (**figs S2F, S3C**). An AlphaFold2 prediction (*25*) of CENP-QU^Foot^ generated a four *α*-helix bundle with high confidence that fitted perfectly into a 3D class of CCAN^ΔCT^ featuring corresponding map density, which we subsequently experimentally validated with a structure of full CCAN bound to DNA.

### Reconstitution of CCAN-CENP-A^Nuc^ onto a 171 bp α-satellite repeat

To assemble a complete CCAN^ΔT^-*α*-satellite nucleosome complex, we reconstituted CCAN^ΔT^ complexes with CENP-C^N^ and CENP-A^Nuc^ using 171 bp *α*-satellite repeat DNA (CENP-A^Nuc^-171). The *α*-satellite DNA exactly matches an *α*-satellite repeat from human chromosome 2, and also shares 93% sequence identity with at least 14 other human centromeres (*26, 27*) (**fig. S4A**). We used two separate 171 bp *α*-satellite sequences, differing in register by 22 bp (termed AS1 and AS2) (**fig. S4A**). To define the exact nucleosome positioning of the AS2 sequence, we determined a cryo-EM structure of a CENP-A^Nuc^-CENP-C^N^ complex at 2.4 Å resolution. From this the nucleotide sequence of the CENP-A^Nuc^ AS2 DNA could be unambiguously assigned (**fig. S4B-D and Table S1**). The dyad axis of the nucleosome is equivalent to NCPs comprising X-chromosome *α*-satellite DNA, with which it shares 71% sequence identity (*11, 27, 28*) (**fig. S4A**). This also matches nucleosome positions defined by *in vivo* ChIP micrococcal nuclease (MNase)-seq analyses of human centromeric chromatin (*16, 28*). Substantial conformational heterogeneity of the DNA termini of CENP-A^Nuc^ is apparent from a 3D variability analysis of the cryo-EM data (**fig. S4E and Movie S1**). Relative to histone H3 nucleosomes (H3^Nuc^), CENP-A^Nuc^ exhibits continuous conformational flexibility of its terminal DNA segments which fluctuate at both ends between being fully wrapped (25% of particles) and unwrapped by approximately 22 bp (also 25% of particles) with the majority of particles being in intermediate states. Unwrapped and flexible terminal DNA segments have previously been observed in yeast and vertebrate CENP-A^Nuc^ structures (*29*), and this correlates with a 110-120 bp nuclease-resistant DNA core (*23, 29–34*), and SAXS data indicating a more open structure for budding yeast CENP-A^Nuc^ (*29*). These *in vitro* studies are also in concordance with *in vivo* ChIP-MNase-seq of CENP-A nucleosomes embedded in natural human centromeres and neocentromeres (*16, 28*). A structure of CENP-A^Nuc^ reconstituted with AS1 (from the CCAN^ΔT^-CENP-A^Nuc^-AS1 cryo-EM data set) was positioned identically to CENP-A^Nuc^-AS2, except the 3’ unwrapped end is absent due the 22 bp 5’ register shift relative to AS2 (**fig. S4A**). In addition to resolving the nucleosomal DNA and histone proteins, we were able to build the CENP-C central domain interacting with the C-terminus of CENP-A, and histones H2A/H2B, a structural feature of CENP-A^Nuc^ recognition conserved from yeast to human (*11, 12, 23, 35*).

### The positively charged CCAN channel grips linker α-satellite DNA of CENP-A^Nuc^

We then determined cryo-EM structures of CCAN^ΔT^-CENP-A^Nuc^ complexes using both the AS1 and AS2 DNA. These two structures were identical except for their different DNA boundaries (**figs S5, S6**). Because the AS1 sequence produced a more stable complex, as judged by SEC, a larger cryo-EM data set was collected with this DNA (**Table S1**).

Cryo-EM micrographs of the CCAN^ΔT^-CENP-A^Nuc^ data sets and 2D-class averages showed particles corresponding to three species: (i) CENP-A^Nuc^-CENP-C^N^, (ii) CCAN^ΔT^ bound to free DNA and (iii) CCAN^ΔT^ bound to CENP-A^Nuc^ (**fig. S5A-C**). Our cryo-EM data showing one CCAN^ΔT^ associated with CENP-A^Nuc^ is in agreement with an assessment of the oligomeric state of CCAN^ΔT^-CENP-A^Nuc^ in solution using SEC-MALS, indicating a single CCAN^ΔT^ assembled onto CENP-A^Nuc^ (**fig. S5D**).

In the CCAN^ΔT^-DNA complex, determined at 4.5 Å resolution, EM density is clearly visible for a linear ∼24 bp DNA duplex threading through the CENP-LN channel (**Figs 2A, 3A and fig. S5E**). The CCAN^ΔT^-CENP-A^Nuc^ complexes exhibited conformational heterogeneity, limiting the resolution of a consensus cryo-EM reconstruction to 8.9 Å (**Fig. 2A and figs S5, S6**). Focused 3D classification and refinement of the CCAN^ΔT^-DNA component alone produced a 7.3 Å resolution reconstruction with a protein and DNA structure identical to that of the CCAN^ΔT^-DNA complex (**fig. S5C, E**). Therefore, we generated a composite CCAN^ΔT^-CENP-A^Nuc^ structure from the higher resolution CENP-A^Nuc^-CENP-C^N^ and CCAN^ΔT^-DNA reconstructions (**Fig. 2A**). In the CCAN^ΔT^-CENP-A^Nuc^ complexes, the CENP-A^Nuc^ DNA is wrapped similarly to the free CENP-A^Nuc^-CENP-C^N^ (**fig. S6**).

**Figure 2.**
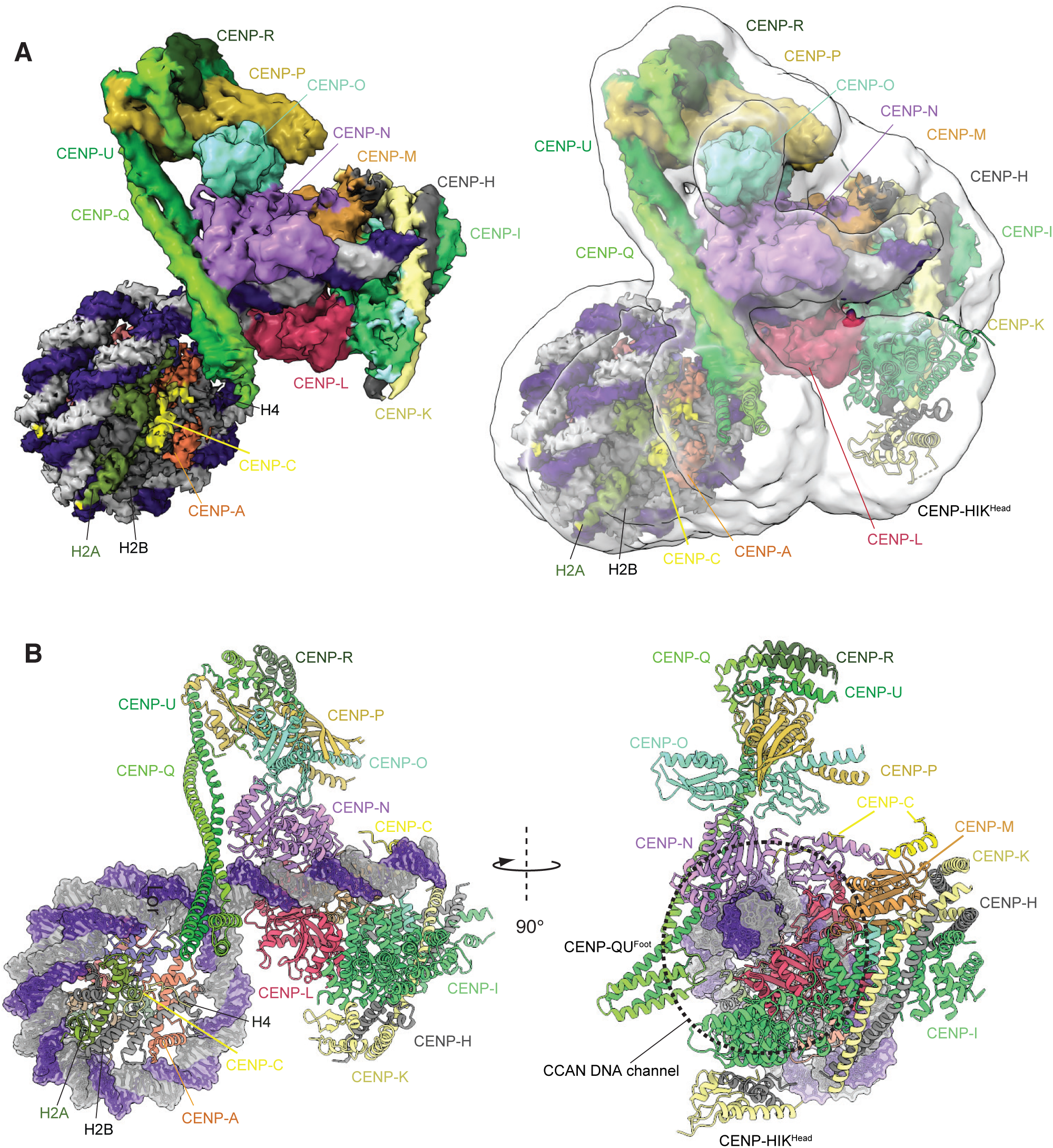
Structure of the CCAN^ΔT^-CENP-A^Nuc^ complex. (**A**) Right panel shows the consensus CCAN^ΔT^-CENP-A^Nuc^ cryo-EM map (transparent white) overlaid onto the composite CCAN^ΔT^-CENP-A^Nuc^ cryo-EM density map based on individual cryo-EM maps for the CENP-A^Nuc^-CENP-C^N^ and CCAN^ΔT^-DNA reconstructions. The left panel shows the composite map alone. (**B**) Two orthogonal views of the CCAN^ΔT^-CENP-A^Nuc^ complex depicted in cartoon representation for protein and space filling for DNA.

**Figure 3.**
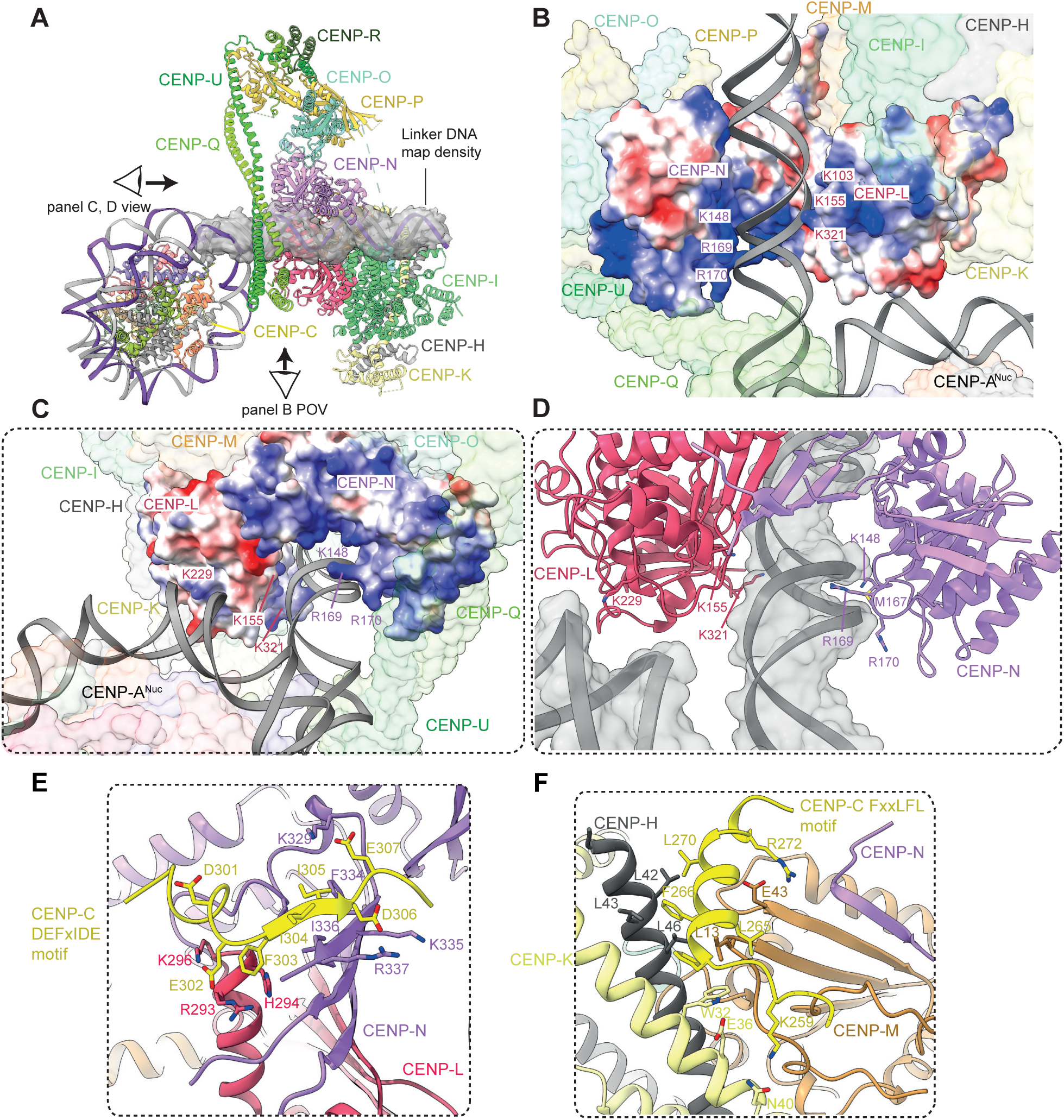
The extranucleosomal linker DNA is gripped by the CENP-LN channel. (**A**) Cartoon representation of CCAN^ΔT^-CENP-A^Nuc^ with cryo-EM density shown for extranucleosomal linker DNA (from CCAN^ΔT^-DNA cryo-EM density map). (**B**) The CENP-LN module features a positively-charged channel that complements the size and charge of a DNA duplex. Electrostatic surface charge shown for CENP-LN. (**C** and **D**). Details of the CENP-LN channel-DNA interaction. R169 of CENP-N inserts into the DNA minor groove. CENP-LN surface charge shown in C. (**E** and **F**) CENP-C binds to two sites on CCAN. The DEFxIDE motif binds to CENP-LN (E), whereas the FxxLFL motif binds to CENP-HIKM (F).

A striking feature of the CCAN^ΔT^-CENP-A^Nuc^ complex is that CCAN^ΔT^ forms few direct contacts with CENP-A^Nuc^, but instead its primary contact with the CENP-A^Nuc^ is through the extended extranucleosomal linker DNA. CENP-A^Nuc^ forms a small end-on contact, mediated through its DNA gyre, with the back-face of CCAN^ΔT^, interacting with a conserved basic surface of CENP-L (**Fig. 3C**). This positions the extranucleosomal DNA to thread through the CENP-LN channel. Clearly resolved cryo-EM density for the DNA phosphate backbone shows it tightly gripped by conserved basic residues of the CENP-LN channel (**fig. S5E**). These create a dense, continuous positively-charged surface complementing the shape and charge of the 24 bp linker DNA duplex (**Figs 2, 3**). Interactions of the side chains of CENP-N residues K148, M167 and R169 with the DNA minor groove contribute to defining a fixed DNA register (**Fig. 3C, D**). In the DNA-bound state, the CENP-LN channel contracts, tightening its grip on the DNA duplex (**fig. S5F**), and the channel is extended by the apposition of CENP-HIK^Head^ in the raised conformation (**Fig. 3A**). The CENP-QU^Foot^ repositions slightly towards CENP-A^Nuc^ and, at low map contour levels, diffuse density bridges CENP-QU^Foot^ with one side of CENP-A^Nuc^ (**Fig. 2A**), possibly originating from the N-terminus of CENP-QU, CENP-C^N^ and the C-terminus of histone H2A.

### CENP-C interaction sites on CENP-LN and CENP-HIKM submodules

The reconstituted CCAN^ΔT^-CENP-A^Nuc^ complex includes CENP-C^N^. The CCAN-interacting region of CENP-C maps to the PEST sequence (CENP-C^PEST^), a 200-residue intrinsically disordered region N-terminal of the CENP-C central domain that features two highly conserved short linear sequence motifs (*8, 19, 20*). Cryo-EM maps of the CCAN^ΔT^-DNA and CCAN^ΔT^-CENP-A^Nuc^ complexes indicated two volumes of unassigned map density not present in apo-CCAN^ΔCT^ and CCAN^ΔC^-DNA complexes (discussed below), associated with the CENP-HIKM and CENP-LN modules (**fig. S6D**), previously shown to interact with CENP-C (*8, 18–20*). To guide the interpretation of these densities, we used AlphaFold2 (*36*) to predict potential models of how CENP-C^PEST^ would interact individually with CENP-LN, and CENP-HIKM. The first run predicted the conserved DEFxIDE motif (residues 301-307) (*8, 18–20*) of CENP-C^PEST^ would form an edge *β*-strand with the CENP-N *β*-sheet (**Fig. 3E**). This position corresponded well to additional cryo-EM density associated with CENP-N (**fig. S6D**). Phe and Ile residues of the DEFxIDE motif dock into a hydrophobic pocket at the CENP-LN interface, whereas the flanking acidic residues form electrostatic interactions with conserved Arg and Lys residues of CENP-N, and CENP-L. In support of this model, a prior study mutating the EFxID residues of the CENP-C DEFxIDE motif abolished its interaction with CENP-LN *in vitro* and disrupted CENP-N recruitment to centromeres *in vivo* without perturbing CENP-C localization (*8*). For CENP-HIKM, AlphaFold2 also predicted that the CENP-C^PEST^ FxxLFL motif (residues 262-267) (*19*) would bind as an *α*-helix to a hydrophobic site at the combined CENP-H-CENP-K-CENP-M interface, involving the conserved Trp32 of CENP-K (**Fig. 3F**). This prediction was validated by cryo-EM density at this location (**fig. S6D**), and is consistent with a previous report that mutating the LFL residues of this motif disrupts CENP-C interactions with CENP-HIKM, *in vitro* and *in vivo* (*19*). Both the LN and HIKM interaction sites are located on the back-face of CCAN, facing the CENP-A^Nuc^, suggesting a mechanism for how CENP-C tethers kinetochores to CENP-A^Nuc^ (*7*).

### α-Satellite linker DNA is a crucial determinant of stable CCAN association with CENP-A^Nuc^

Our CCAN^ΔT^-CENP-A^Nuc^ structure suggests that the interaction of the extranucleosomal DNA duplex with the CENP-LN channel is a major determinant of CCAN assembly onto a centromeric *α*-satellite-CENP-A^Nuc^. To test this, we performed reconstitutions assessing the role of the extranucleosomal DNA, CENP-C^N^, basic residues lining the CENP-LN channel, CENP-HIK^Head^ and CENP-QU^Foot^. We first tested the requirement of the extranucleosomal DNA. Using SEC, we found that CENP-A^Nuc^ wrapped with 147 bp of *α*-satellite DNA (*11, 27, 37*) did not form stable complexes with CCAN^ΔT^ in the absence of CENP-C^N^, whereas a 171 bp *α*-satellite that included the extranucleosomal DNA formed stable complexes with CCAN^ΔT^ with and without CENP-C^N^ (**fig. S7A, B**). Interactions with CENP-A^Nuc^ reconstituted with a 171 bp *α*-satellite were equally stable in the presence or absence of the B-box, consistent with sequence-independent interactions with the DNA phosphate backbone (**fig. S7C**).

Having established a requirement for linker DNA in the assembly of CCAN^ΔCT^-CENP-A^Nuc^ complexes, we next assessed how modifications to the CENP-LN channel affected CCAN^ΔT^ interactions with CENP-A^Nuc^. In the absence of CENP-C^N^, mutating three positive patches of CENP-L (CENP-Lcm), that interact with linker DNA and the nucleosome gyre, specifically abolished binding of CCAN^ΔCT^ to CENP-A^Nuc^-171 (**Fig. 3 and figs S5F, S7D**), without perturbing CCAN assembly. Identical results were obtained by deleting either CENP-HIK^Head^ or CENP-QU^Foot^, consistent with our cryo-EM structure, although deleting CENP-QU^Foot^ also resulted in defects in CCAN assembly (**fig. S8A, B**). These deletions also moderately reduced CCAN binding to isolated DNA (**fig. S8C**). To assess the validity of the CCAN-CENP-A^Nuc^ model in a cellular context, we tested the effect of the CENP-Lcm mutant that disrupted CCAN^ΔCT^ binding to CENP-A^Nuc^-171 *in vitro*, on centromere localization of CENP-L *in vivo*. In HEK293 cells, localization of the CENP-Lcm mutant to kinetochores was substantially impaired as compared to wild-type CENP-L (**fig. S8D**).

In previous structures of the CENP-N N-terminal domain (CENP-N^NT^) associated with CENP-A^Nuc^ (*8–11*), CENP-N^NT^ interacts with the L1 loop of CENP-A (CENP-A^L1 loop^) and adjacent DNA gyre at SHL2-3. In agreement with these structures, we found that mutating CENP-A^L1 loop^ disrupts the interaction of the isolated CENP-LN dimer with CENP-A^Nuc^ wrapped with 171bp *α*-satellite DNA (**fig. S9A).** By contrast, our structure of CCAN-CENP-A^Nuc^-171 does not involve an interaction of CENP-N^NT^ with CENP-A^L1 loop^, and consistent with this, mutating CENP-A^L1 loop^ did not disrupt interactions of CENP-A^Nuc^-171 with CCAN^ΔCT^ as assessed by both pull-down assays and during reconstitution on SEC (**fig. S9A, B**). These results, as well as previous structural information (*8–11*), suggest that the mode of interaction between CENP-LN and CENP-A^Nuc^ is different when CENP-LN is part of the full CCAN complex compared to CENP-LN alone. To further test this hypothesis, we performed competition assays where we added the CENP-N^NT^, which binds specifically to CENP-A^L1 loop^, at a four-fold excess over the CCAN-CENP-A^Nuc^ complex (**fig. S9C**). We observed that the CCAN-CENP-A^Nuc^ complex readily accommodated the additional CENP-N^NT^, without evident diminution of CCAN binding, and this additional CENP-N^NT^ incorporation was dependent on CENP-A^L1 loop^. This suggests that in a fully assembled CCAN-CENP-A^Nuc^ complex, CENP-A^L1 loop^ is not occupied and therefore available for CENP-N^NT^ binding (**fig. S9D**). These biochemical experiments are all consistent with, and support our cryo-EM reconstruction. Modelling a CENP-N^NT^-based interaction of CCAN with CENP-A^L1 loop^ shows that CENP-A^Nuc^ would clash extensively with the CENP-L, CENP-HIK and CENP-OPQUR modules (as well as CENP-TWSX module as discussed below) (**fig. S9E**). In both models, CENP-N^NT^ forms the same interactions with the DNA backbone, although at SHL7-8 for CENP-N^NT^ in the context of CCAN (**fig. S9F**). Thus, previously described mutants at the CENP-N^NT^-DNA interface that disrupt CENP-N-centromere localization *in vivo* (*6, 8, 9*), are consistent with, but cannot distinguish between, both modes of CENP-N-CENP-A^Nuc^ interaction. Overall, it appears that the interaction of free CENP-LN with CENP-A^Nuc^ is strictly dependent on CENP-A^L1 loop^ but completely independent of the linker DNA. On the other hand, the interaction between CCAN and CENP-A^Nuc^ is dependent on the linker DNA but not on CENP-A^L1 loop^, consistent with a different mode of interaction between CENP-LN and CENP-A^Nuc^ when CENP-LN is part of the full CCAN, and a previous observation that the L1 loop is not required for the *in vitro* assembly of a functional kinetochore (*38*).

### CENP-C confers selectivity of assembled CCAN for CENP-A mono-nucleosomes

The only direct interaction between the fully assembled CCAN^ΔT^ and the CENP-A histone we observed in our structure is mediated by CENP-C^N^. To validate this finding biochemically, we produced human H3 nucleosomes reconstituted either with a 147 bp 601 sequence or a 601 sequence extended at the 5’ end by 30 bp (177 bp) (**fig. S4A**). As expected from our structure, CCAN^ΔT^ failed to bind to H3^Nuc^ reconstituted with the minimal 147 bp 601 sequence, whereas CCAN^ΔT^ bound to CENP-A^Nuc^ with the same DNA only in the presence of CENP-C^N^, similar to when 147 bp *α*-satellite DNA was used (**figs S7A, and S10A**). Consistent with our structure, binding was observed with the extended 177 bp sequence for both H3^Nuc^-177 and CENP-A^Nuc^-177 (**fig. S10B**). The binding of CCAN^ΔCT^ to CENP-A^Nuc^-177 but not to H3^Nuc^-177, was enhanced 4-5 fold by CENP-C^N^, consistent with selective recognition of CENP-A^Nuc^ by CENP-C (**fig. S10C**). A near identical degree of specificity for CENP-A^Nuc^ compared to H3^Nuc^ reconstituted with the 147 bp 601 sequence was observed previously for the CENP-CHIKMLN sub-complex (*39*).

### CENP-TWSX binds and partially wraps linker DNA

The CCAN^ΔT^-CENP-A^Nuc^ structure we describe was reconstituted without the CENP-TWSX module due to our initial difficulties incorporating CENP-TWSX into CCAN, similar to other reports (*40*). We subsequently found that removing all affinity tags from CENP-WSX enabled CENP-TWSX incorporation into CCAN (**fig. S10D**), and as judged by SEC, this 16-subunit holocomplex interacts with the 171 bp *α*-satellite-CENP-A^Nuc^ similarly to CCAN^ΔT^ (**fig. S10D**), consistent with CENP-A, CENP-C and CENP-T forming a single 20-subunit inner kinetochore complex (*41*). SEC experiments showed that the full CCAN complex also requires extranucleosomal linker DNA to bind CENP-A^Nuc^ in the absence of CENP-C, suggesting that CENP-TWSX does not change the fundamental mechanism of CCAN assembly onto CENP-A^Nuc^ (**fig. S10E**). EMSA experiments revealed that CENP-TWSX increases the affinity of CCAN^ΔCT^ for both CENP-A^Nuc^ and H3^Nuc^ nucleosomes containing 177 bp 601 DNA approximately 3-fold, suggesting that it does not confer additional selectivity for CENP-A^Nuc^ over canonical H3^Nuc^ (**fig. S10 C, F**).

To understand how CENP-TWSX contributes to CCAN recognition of DNA and CENP-A^Nuc^, we determined cryo-EM structures of CCAN^ΔC^ with a 53 bp DNA linker segment (**fig. S4A**), and also the complete 16-subunit CCAN in complex with CENP-A^Nuc^. In the presence of DNA, full CCAN becomes rigidified which allowed us to obtain a well-resolved cryo-EM map at 2.8 Å resolution with all regions of CCAN clearly defined (**Fig. 4A, B, fig. S11, Table S1 and Movie S2**). The four histone fold domains of CENP-TWSX resemble the histone H3-H4 tetramer, and are similar to crystal structures of isolated chicken CENP-TWSX (*42*). CENP-TWSX forms multiple interactions with neighboring CCAN subunits and DNA (**Fig. 4B, C**). Contacts with a modestly repositioned CENP-HIK^Head^ are mediated by both CENP-T as well as the extended N terminus of CENP-W (*43, 44*). CENP-W also directly contacts the CENP-N pyrin domain. Additionally, CENP-TWSX bridges CENP-HIK^Head^ and CENP-QU^Foot^ through an interaction involving CENP-Q and CENP-S. The CENP-TWSX-mediated linkage of CENP-HIK^Head^ to CENP-N and CENP-QU^Foot^ creates an enclosed chamber that topologically entraps the extranucleosomal DNA (**Fig. 4A-D**). The complete chamber thus comprises CENP-LN, CENP-HIK and CENP-TW modules but is augmented and rigidified by all other CCAN subunits. As in the CCAN^ΔT^-DNA complex, the DNA duplex inserts into the deep CENP-LN channel, however the presence of CENP-TWSX and repositioned CENP-HIK^Head^, induces a marked curvature of the DNA. Strikingly, the DNA starts to wrap around the CENP-TW histone fold domains as it emerges from the CENP-LN channel in a manner comparable to how a canonical H3-H4 tetramer wraps DNA (**Fig. 4E**). However, unlike the H3-H4 tetramer, this DNA binding is also supported and completed by CENP-HIK^Head^. This results in the DNA duplex engaging with a 50 Å-long positively-charged groove at the interface of CENP-I^Head^ and CENP-TW (**Fig. 4D**). Also similar to nucleosomes, the N-terminus of the CENP-T (HFD) inserts into the minor groove of the extranucleosomal DNA positioning the highly conserved Arg450 to contact the DNA backbone (**Fig. 4E**). The CENP-LN channel and CENP-TW-CENP-I^Head^ groove perfectly complement the shape and charge of the DNA duplex, with the DNA phosphate backbone forming a continuous interface with CCAN along its entire 40 bp length, burying some 976 Å^2^ of DNA surface area.

**Figure 4.**
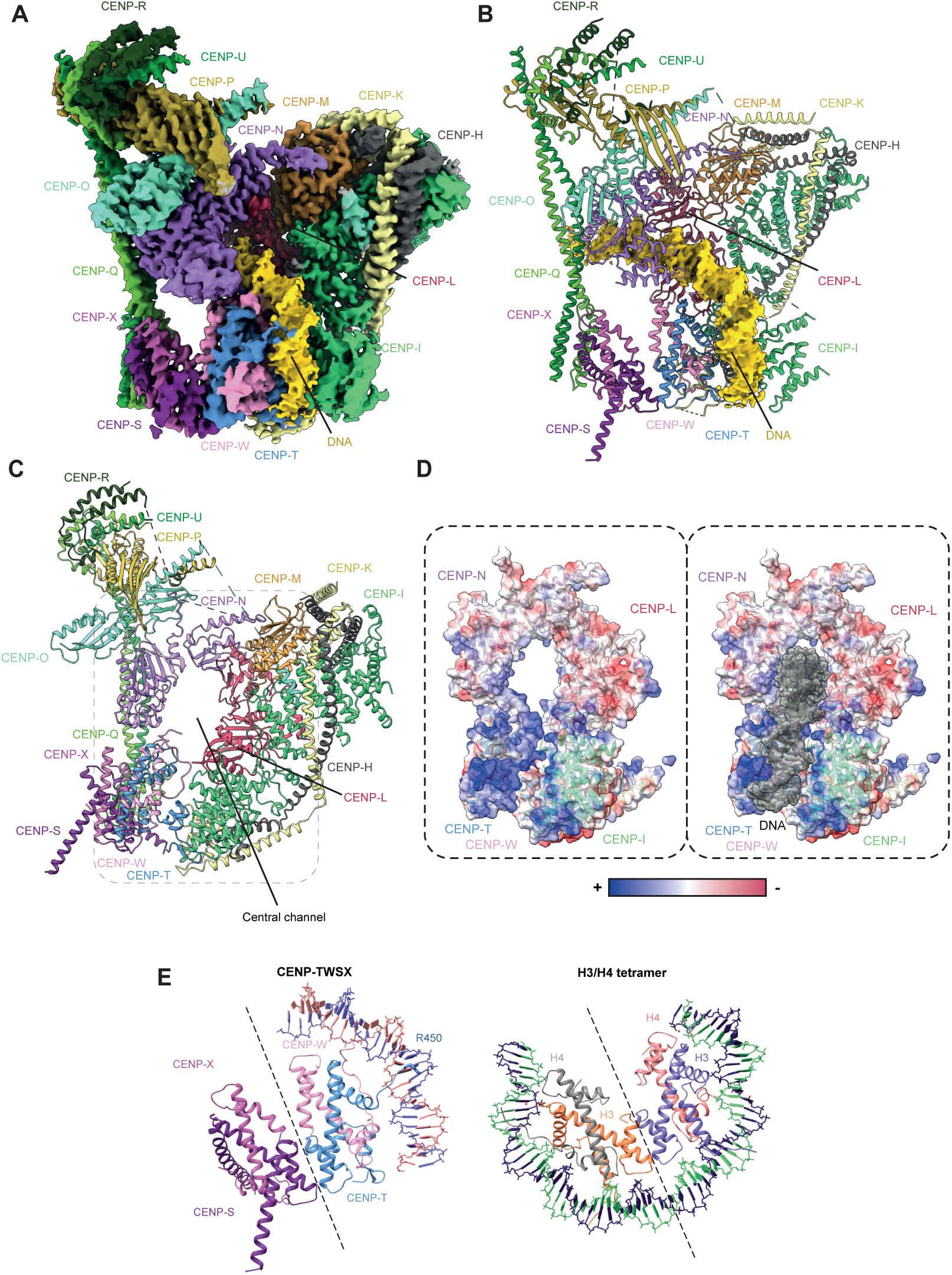
Architecture of the full CCAN^ΔC^-DNA complex. (**A**) Cryo-EM density map, and (**B**) molecular model of the CCAN^ΔC^-DNA complex with DNA map density shown in yellow. (**C**) Molecular organization of CCAN generates a completely enclosed, narrow chamber that can topologically entrap linker DNA. (**D**) Electrostatic representation of the central DNA-binding chamber of the CCAN formed from the CENP-LN, CENP-TW and CENP-HIK^Head^ modules shows a highly positively charged chamber that tightly grips the DNA. (**E**) The CENP-TWSX module strongly resembles the canonical H3/H4 nucleosome tetramer. However, CENP-TWSX only partially wraps the DNA along its surface compared to the nucleosome module.

To complete our structural analysis, we determined cryo-EM structures of the 16-subunit CCAN in complex with CENP-A^Nuc^ reconstituted with 171 bp *α*-satellite DNA (**Fig. 5A, B and fig. S11C**). CCAN assembles onto CENP-A^Nuc^ similarly to CCAN^ΔT^. The nucleosome associates with the back-face of CCAN, and the extranucleosomal DNA threads through the CENP-LN channel. The linker DNA of the *α*-satellite repeat is gently curved and interacts with CENP-TW-CENP-I^Head^ reminiscent of the CCAN^ΔC^-DNA complex. We note that the AS2 *α*-satellite DNA of 171 bp in length is approximately 15 bp too short to wrap all the way around the CENP-TWSX module.

**Figure 5.**
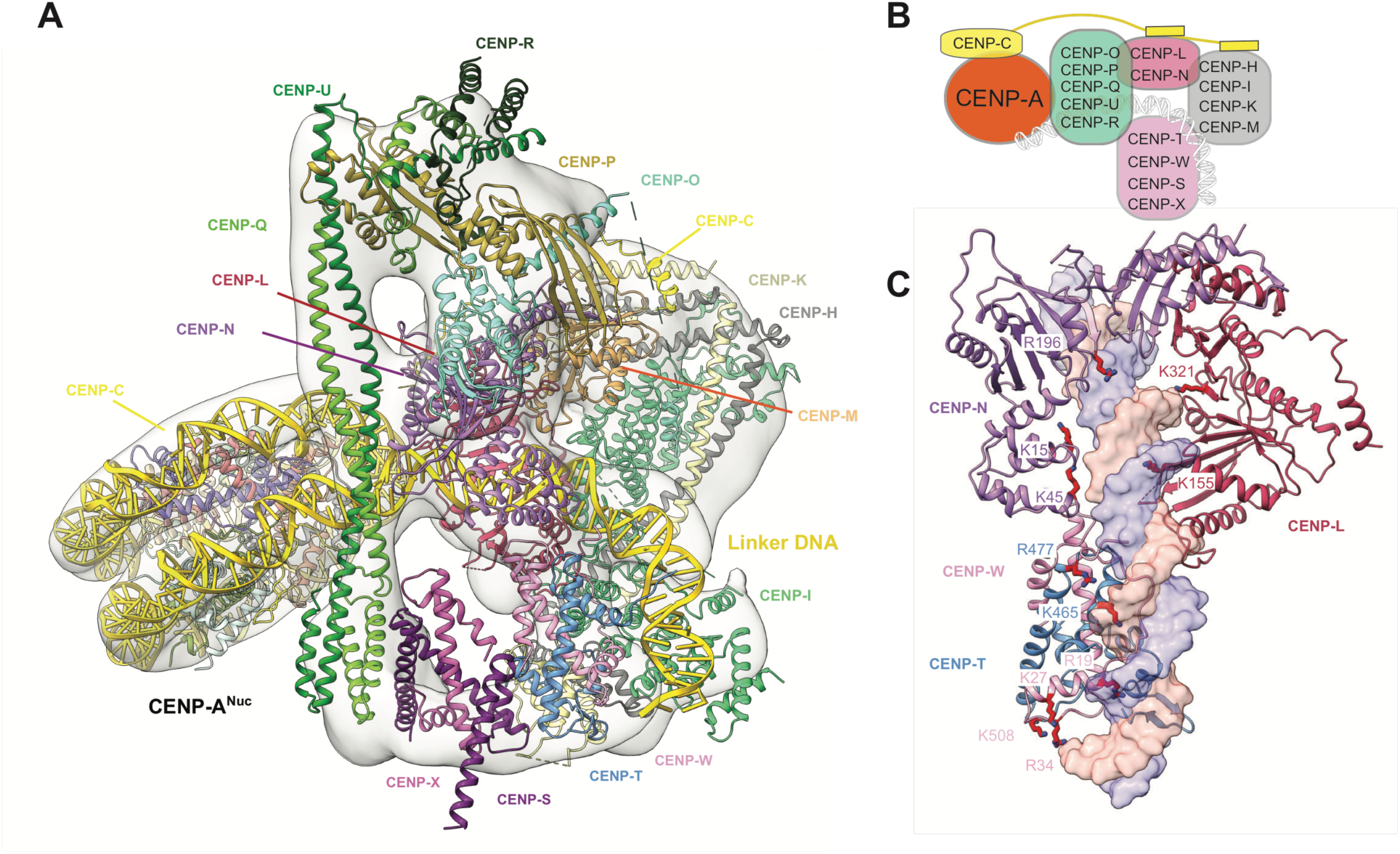
Structure of the complete CCAN-CENP-A inner kinetochore module. **(A**) Complete atomic model of the CCAN-CENP-A^Nuc^ complex fitted into the low-resolution cryo-EM reconstruction of CCAN-CENP-A^Nuc^ complex determined using AS2 *α*-satellite repeat. **(B)** A cartoon schematic of the CCAN-CENP-A^Nuc^ complex. (**C**) Mutations of CENP-N and CENP-TW that were previously shown to impair centromere function contribute to the central DNA-binding channel of the CCAN (*6, 8–10*). Residues implicated in DNA binding by CENP-L, and mutated in the current study, are also shown.

We observed that CENP-TW contributes to DNA binding via conserved basic residues that were previously implicated in mediating DNA binding *in vitro*, and for CENP-W, are required for mitotic progression, and its recruitment to kinetochores *in vivo*. These functions of CENP-TW, and CENP-TWSX’s ability to supercoil DNA, lead to proposals that CENP-TWSX is a DNA-binding nucleosome-like particle, consistent with the curvature of the DNA in our structure (*42, 45*). These residues of CENP-TW, together with basic residues of CENP-N, whose mutation disrupt CENP-N association with CENP-A^Nuc^ *in vitro*, and centromere localization *in vivo* (*6, 8–10*), form a continuous spine lining the conserved CCAN-DNA interface (**Fig. 5C**).

## Discussion

Our study of the human CCAN-CENP-A^Nuc^ complex defines the architecture and assembly of this foundational module of the kinetochore, and reveals how CCAN utilizes *α*-satellite linker DNA to stabilize its assembly on nucleosomes. We show that nucleosomal DNA leads into a continuous linker DNA tightly gripped by the CENP-LN channel, and clamped from below by the CENP-TW-CENP-I^Head^ platform. The remarkably similar features of DNA duplex recognition by the basic CENP-LN channel of *Hs*CCAN and *Sc*CCAN (*23*), suggests a common evolutionary origin of centromere-kinetochore assembly. In humans, however, the DNA duplex engaged by CCAN is the *α*-satellite linker DNA rather than the unwrapped CENP-A^Nuc^ DNA of budding yeast, and *Hs*CCAN entraps the centromeric DNA inside an enclosed chamber formed from CENP-LN and CENP-TW-CENP-HIK^Head^. CCAN can engage ∼40 bp of extranucleosomal DNA, which could be facilitated either by unwrapping of ∼16 bp of the upstream CENP-A^Nuc^, consistent our structures showing a continuum of DNA termini wrapping by CENP-A^Nuc^, and/or by the sparse occupancy of CENP-A^Nuc^ on centromeric DNA (*46*). Such organization is consistent with evidence that CENP-C and CENP-T bridge adjacent nucleosomes (*41*). Crucially, we show here that CCAN tightly grips and topologically entraps the linker DNA, providing a molecular explanation for how the inner kinetochore can withstand strong pushing and pulling forces applied by the mitotic spindle from any direction during mitosis, a central function of this large macromolecular assembly.

Our finding that CCAN can bind to either canonical or CENP-A nucleosomes bearing adequate linker DNA could provide an explanation for how the inner kinetochore maintains attachments at centromeres where H3^Nuc^ vastly exceeds CENP-A^Nuc^ across the entire length of centromeric *α*-satellite repeats (*47*). Furthermore, it could explain why synthetic depletion of CENP-A once CCAN is assembled onto centromeres, nevertheless permits mitotic centromere function in human cells (*48*), as fully assembled CCAN could fulfil its functions by capturing linker DNA. Salt extraction of CENP-A did not reduce CENP-N and CENP-C levels in centromeric chromatin (*32*), providing evidence that once CCAN is assembled at centromeres in a CENP-A dependent process (*49, 50*), it no longer requires CENP-A for centromere localization. Lastly, our structure also provides a possible explanation for how some organisms that lack CENP-A yet still depend on CCAN for chromosome segregation might function, as CCAN bound to the linker DNA in those organisms could be sufficient to couple chromosomes to the mitotic spindle (*51*).

An emerging question from this study is how human CCAN localizes almost exclusively at the centromere given its modest preference for CENP-A^Nuc^ mono-nucleosomes compared to H3^Nuc^ (*47*). One possible explanation is that the CENP-LN module is selectively recruited to the centromere, through recognition of CENP-A^L1 loop^, during early stages of the cell cycle, and is then displaced from the L1 loop to form a fully assembled CCAN, thus confining kinetochore assembly to a specific chromatin locus. CDK-mediated CENP-C phosphorylation phosphorylation (*12, 18, 52*) and/or chromatin compaction (*17, 18*) might regulate such displacement. CENP-B recognizes the B-box motifs of the *α*-satellite repeat and interacts with CENP-C, providing another possible mechanism for specific CCAN recruitment (*53*). However, B-box motifs are absent from some neo-centromeres and the Y-chromosome yet CCAN recruitment is still specific (*54*). Different degrees of wrapping of CENP-A^Nuc^ in centromeric arrays compared to H3^Nuc^ might also provide selectivity for CCAN assembly, as could the H1 histone that would block access of CCAN to the linker DNA at H3^Nuc^ (*55*). The architecture of CCAN within centromeric arrays of regional kinetochores is an outstanding question.

This study also highlights the common mechanisms of topological entrapment used by biological systems responsible for accurately mediating chromosome segregation, such as the cohesin complex that, likewise resisting vast forces exerted by the spindle, maintains cohesion between sister chromatids.

## Acknowledgments

We are grateful to the LMB and eBIC EM facilities, and Diamond Light Source for help with the EM and X-ray data collection, J. Grimmett and T. Darling for computing, A. Burt for assistance with 3D variability analysis and J. Shi for help with insect cell expression.

## Funding

UKRI/Medical Research Council MC_UP_1201/6 (DB)

Cancer Research UK C576/A14109 (DB)

Boehringer Ingleheim Fonds Fellowship (SY)

## Author contributions

D.B., K.W.M. and S.Y. designed the study and experiments. K.W.M. and S.Y. cloned constructs except H3 nucleosome, which was cloned by Z.Z. Z.Z. produced and purified all DNA substrates. S.Y. and K.W.M. purified all proteins with help from J.Y., Z.Z. and M.P. S.Y. and K.W.M. performed all biochemical experiments, prepared EM grids, collected and processed the cryo-EM data and built atomic models together with D.B. T.D. helped with cryo-EM data processing. T.T. performed all *in vivo* experiments. Dom B. crystallized OPQUR^Foot^ and collected crystallography data and assisted with crystallography data processing. S.H.M. performed SEC-MALS experiments. D.B., K.W.M. and S.Y. wrote the manuscript with input from all authors.

## Competing interests

Authors declare that they have no competing interests.

## Data and materials availability

PDB and cryo-EM maps have been deposited with RCSB and EMDB, respectively. Accession numbers are listed in Tables S1 and S2.

## Materials and Methods

### Cloning of the CCAN and CENP-A protein complexes

Genes encoding CENP-O, CENP-P, CENP-Q, CENP-U, CENP-R, CENP-L, CENP-N, CENP-T, CENP-W, CENP-S, CENP-X were synthesized by Thermo Fisher Scientific with codon optimization for *Trichoplusia ni*. Genes for the CENP-OPQUR and CENP-TWSX modules were subsequently cloned separately into a modified Multibac expression system (*56*). TEV cleavable double strep II (DS) tags were added to the C-termini of CENP-U and CENP-P. An N-terminal 6xHis-SNAP tag was added to CENP-T. CENP-L and CENP-N were sub-cloned into a pF1 plasmid with C-terminal 3C-cleavable SNAP-DS tags on CENP-N and an N-terminal 3C-cleavable GST-tag on CENP-L. Genes for CENP-C, CENP-H, CENP-I, CENP-K and CENP-M were cloned from human cDNA. Genes for the CENP-HIKM module were sub-cloned into a pF1 plasmid (*56*) with a C-terminal TEV-cleavable DS tag on CENP-I. The N-terminal fragment of CENP-C (residues 1-544) (CENP-C^N^) was cloned into a modified pFastBac vector with a C-terminal 3C-cleavable SNAP-DS tag.

CENP-LN^cm^ (R46D, R47D, K48E, K155E, K321E mutated in CENP-L), CENP-LN^bs^ (R236, I237, I238, H239, E240 of CENP-N mutated to alanine), CENP-O^Δ35^PQUR (CENP-PQUR^full-length^, CENP-O^Δ35^), CENP-HIKM^ΔHead^ (CENP-H^1-183^, CENP-I^340-756^, CenpM^full-length^, HsCENP-K^1-151^), CENP-OPQUR^Foot^ (CENP-OPR^full-length^, CENP-Q^133-268^, CENP-U^295-418^), CENP-OPQUR^ΔN^ (CENP-OPR^full-length^, CENP-Q^79-268^, CENP-U^250-418^) were generated using a HiFi DNA assembly kit (New England Biolabs).

For CENP-HIK^Head^ expression, DNA encoding CENP-I 1-280, CENP-H 204-247, and CENP-K 165-269 were cloned, in that order, as a single polycistron into pRsfDuet using HiFi assembly (New England Biolabs), with a 3C-SNAP-2xStrepII tag in frame with CENP-I, thereby yielding the pRsfDuet CENP-HIK^Head^ expression vector.

The N-terminal fragment of CENP-N (residues 1-212) was cloned into pGex6P-1 in-frame with an N-terminal 3C-cleavable GST tag using HiFi assembly (New England Biolabs, USA).

Human cDNAs comprising CENP-A and H4 were cloned by restriction-ligation into ORF1 and ORF2 of pRsfDuet. A bicistronic cassette encompassing 6xHis3C-tagged H2A and untagged H2B was cloned by restriction-ligation into pAcycDuet.

The coding regions of histone H2A, H2B, H4 and H3 were amplified by PCR from cDNA and cloned into pET28 plasmid. A double Strep-II tag together with a TEV cleavage site was attached to the N-termini of H3 and H2A proteins. For H3 octamer reconstitution, the histones H2A, H2B, H3 and H4 were assembled into a single pET28 plasmid by the USER methodology.

### DNA fragment generation and purification

For DNA fragment preparation, two complementary oligos were synthesized by Sigma-Aldrich. Mixing the oligos in 1x PCR reaction mixture, the fragments were produced by single step extension at 68°C for 1 min. The final products were purified by 1 ml Resource Q anion exchange chromatography (Cytiva) and stored in a buffer of 2 M NaCl, 10 mM Tris.HCl (pH 7.5), 1 mM EDTA, 2 mM DTT at -20°C.

Oligos aSat147F (ATCGAGGAAG TTCATATAAA AGGCAAACGG AAGCATTCTC AGAATATTCT TTGTGATGAT GGAGTTTCAC TCACAGAGCT GAAC) and aSat174R (ATCAAATATC CACCTGCAGA TTCTACCAAA AGTGTATTTG GAAACTGCTC CATCAAAAGG CATGTTCAGC TCTGTGAGTG AAAC) were used for generation of αSat147. Oligs aSat171F(AATCTGCAAG TGGATATTTG GACCGCTTTG AGGCCTTCGT TGGAAACGGG AATATCTTCA CATAAAAACT AAACAGAAGC ATTCTCAGAA ACTTC) and aSat171R (CTACAAAAAG AGTGTTTCAA AACTGCTCTA TCAAAAGGAA TGTTCAACTC TGTGAGTTGA ATGCAATCAT CACAAAGAAG TTTCTGAGAA TGCTTC) were used for AS1 generation. Oligos aSat171BbF (CCGCTTTGAG GCCTTCGTTG GAAACGGGAA TATGTTCACA TAAAAACTAG ACAGAAGCAT TCTCAGAAAC TTCTATGTGA TGTTTGCATT CAACT) and aSat171BbR (TCCAAATGTC CAATTCCAGA TACTACAAAA AGAGTGTTTC AAAACTGCTC TATGAAAAGG AATGTTCAAC TCTATGAGTT GAATGCAAAC ATCACA) were used for AS2 generation. Oligos 171aSatAaF (CCGCTTTGAG GC CTA TGGTGAAAAA GGAA A TATGTTCACA TAAAAACTAG ACAGAAGCAT TCTCAGAAAC TTCTA TGTGA TGTTTGCATT CAACT) and 171aSatAaR (TCCAAATGTC CAATTCCAGA TACTACAAAA AGAGTGTTTC AAAACTGCTC TATGAAAAGG AATGTTCAAC TCTATGAGTT GAATGCAAAC ATCACA) were used for AS2^ΔB-box^ generation. Oligos 177a601F (TTGGACCGCT TTGAGGCCTT CGTTGGAAAC GGGAAGAATC CCGGTGCCGA GGCCGCTCAA TTGGTCGTAG ACAGCTCTAG CACCGCTTAA ACGCACGTAC) and 177a601R (ATCGGATGTA TATATCTGAC ACGTGCCTGG AGACTAGGGA GTAATCCCCT TGGCGGTTAA AACGCGGGGG ACAGCGCGTA CGTGCGTTTA AGCGGTG) were used for 177-601 generation. Oligos widom 601F(ATCGAGAATC CCGGTGCCGA GGCCGCTCAA TTGGTCGTAG ACAGCTCTAG CACCGCTTAA ACG CACGTAC GCGCTGTCCC) and widom601R (ATCGGATGTA TATATCTGAC ACGTGCCTGG AGACTAGGGA GTAATCCCCT TGGCGGTTAA AACGCGGGGG ACAGCGCGTA CGTG) were used for 601 generation. Oligos cy5-601F (cy5-ATCGAGAATC CCGGTGCCGA GGCCGCTCAA TTGGTCGTAG ACAGCTCTAG CACCGCTTAA ACG CACGTAC GCGCTGTCCC) and widom601R (ATCGGATGTA TATATCTGAC ACGTGCCTGG AGACTAGGGA GTAATCCCCT TGGCGGTTAA AACGCGGGGG ACAGCGCGTA CGTG) were used for Cy5-601 generation. 53 bp linker DNA was synthesized by IDT and prepared as double-stranded DNA.

### DNA sequences used in this study

**AS1**: AATCTGCAAGTGGATATTTGGACCGCTTTGAGGCCTTCGTTGGAAACGGGAATATCT TCACATAAAAACTAAACAGAAGCATTCTCAGAAACTTCTTTGTGATGATTGCATTCA ACTCACAGAGTTGAACATTCCTTTTGATAGAGCAGTTTTGAAACACTCTTTTTGTAG.

**AS2**: CCGCTTTGAGGCCTTCGTTGGAAACGGGAATATGTTCACATAAAAACTAGACAGAAGCATTCTCAGAAACTTCTATGTGATGTTTGCATTCAACTCATAGAGTTGAACATTCCTTTTCATAGAGCAGTTTTGAAACACTCTTTTTGTAGTATCTGGAATTGGACATTTGGA.

**AS2^ΔB-box^**: CCGCTTTGAGGCCTATGGTGAAAAAGGAAATATGTTCACATAAAAACTAG ACAGAAGCATTCTCAGAAACTTCTATGTGATGTTTGCATTCAACTCATAG AGTTGAACATTCCTTTTCATAGAGCAGTTTTGAAACACTCTTTTTGTAGT ATCTGGAATTGGACATTTGGA.

**aSat147**: ATCGAGGAAGTTCATATAAAAGGCAAACGGAAGCATTCTCAGAATATTCTTTGTGATGATGGAGTTTCACTCACAGAGCTGAACATGCCTTTTGATGGAGCAGTTTCCAAATACACTTTTGGTAGAATCTGCAGGTGGATATTTGAT.

**601:** ATCGAGAATCCCGGTGCCGAGGCCGCTCAATTGGTCGTAGACAGCTCTAGCACCGCTTAAACGCACGTACGCGCTGTCCCCCGCGTTTTAACCGCCAAGGGGATTACTCCCTAGTCTCCAGGCACGTGTCAGATATATACATCCGAT.

**177-601:** TTGGACCGCTTTGAGGCCTTCGTTGGAAACGGGAAGAATCCCGGTGCCGAGGCCGCTCAATTGGTCGTAGACAGCTCTAGCACCGCTTAAACGCACGTACGCGCTGTCCCCCGCGTTTTAACCGCCAAGGGGATTACTCCCTAGTCTCCAGGCACGTGTCAGATATATACATCCGAT.

**DNA^53^**: AATCTGCAAGTGGATATTTGGACCGCTTTGAGGCCTTCGTTGGAAACGGGAAT.

### Expression and purification of the CCAN and CENP-A protein complexes

The baculoviruses for expression of all CCAN protein complexes were generated using standard protocols (*56*). All the CCAN sub-complexes were expressed individually in High-5 insect cells. The High-5 insect cell line was not tested for mycoplasma contamination and was not authenticated. Typically, 4L of High-5 cells were infected 2.5% v/v with the P3 cell culture and the cells were harvested 48-72 h after infection by centrifugation.

The cells for CENP-OPQUR, CENP-HIKM, CENP-LN, CENP-C^N^ and their associated mutants were lysed with a sonicator in lysis buffer (50 mM Tris.HCl pH 8.0, 300 mM NaCl, 0.5 mM TCEP and 5% glycerol) supplemented with benzamidine, EDTA-free protease inhibitor tablets and benzonase. Clarified lysate was loaded onto a Strep-Tactin column (Qiagen), immobilized proteins washed with lysis buffer, and the complexes were eluted in a buffer containing 300 mM NaCl, 20 mM HEPES pH 7.8, 1 mM TCEP, 5% glycerol with addition 5 mM desthiobiotin (Sigma). For CENP-HIKM, TEV protease was added to remove the DS tag and 3C protease was added to CENP-LN to remove tags.

For the CENP-OPQUR module, the Strep-Tactin eluate was loaded onto the HiTrap SP HP cation exchange column (Cytiva). The column was washed with CENP-OPQUR wash buffer (20 mM HEPES pH 7.8, 300 mM NaCl, 1 mM TCEP, 5% glycerol) supplemented with 10 mM ATP and 10 mM MgCl_2_ for 5 column volumes. The complex was eluted in a single step with 5 column volumes of CENP-OPQUR elution buffer (20 mM HEPES pH 7.8, 500 mM NaCl, 1 mM TCEP, 5% glycerol), aliquoted and flash frozen in liquid nitrogen.

For CENP-OPQUR^Foot^, CENP-OPQUR^ΔN^ and CENP-O^Δ35^PQUR mutants, the Strep-Tactin eluate was diluted to a final salt concentration of 150 mM NaCl directly and loaded onto the HiTrap Heparin HP column (Cytiva). The column was washed with CENP-OPQUR wash buffer supplemented with 10 mM ATP and 10 mM MgCl_2_ for 5 column volumes. The protein was eluted in 20 mM HEPES pH 7.8, 1 mM TCEP and 5% glycerol with NaCl gradient from 150 mM to 600 mM. The peak fraction was flash frozen in liquid nitrogen.

For CENP-HIKM and CENP-HIKM^Head^ sub-complex, the Strep-Tactin eluate was diluted to a final salt concentration of 150 mM NaCl directly and loaded onto the Resource Q anion exchange column (Merck). The protein was eluted in 20mM HEPES pH 8.0, 1 mM TCEP and 5% glycerol with NaCl gradient from 150mM to 600 mM. The peak fraction was concentrated and further purified using a Superdex 200 column (Cytiva) in gel filtration buffer (20 mM HEPES pH 7.8, 300 mM NaCl, 1 mM TCEP). The eluted CENP-HIKM complex was concentrated and flash frozen in liquid nitrogen.

For CENP-C^N^, the Strep-Tactin eluate was loaded directly onto the Resource Q anion exchange column (Cytiva). The protein was eluted in 20 mM HEPES pH 8.0, 1 mM TCEP and 5% glycerol with NaCl gradient from 300 mM to 600 mM. The peak fraction was concentrated and further purified using Superdex 200 column (Cytiva) in gel filtration buffer (20 mM HEPES pH 7.8, 300 mM NaCl, 1 mM TCEP).

For CENP-LN, CENP-LN^cm^ and CENP-LN^bs^ the Strep-Tactin eluate was diluted to a final salt concentration of 150 mM NaCl directly and loaded onto the HiTrap Heparin HP column (Cytiva). The column was washed with CENP-LN wash buffer (20 mM HEPES pH 7.8, 150 mM NaCl, 1 mM TCEP, 5% glycerol) supplemented with 10 mM ATP and 10 mM MgCl_2_ for 5 column volumes. The protein was eluted in 20 mM HEPES pH 7.8, 1 mM TCEP and 5% glycerol with NaCl gradient from 150 mM to 600 mM. The peak fraction was concentrated and further purified using Superdex 200 column (Cytiva) in gel filtration buffer (20 mM HEPES pH 7.8, 300 mM NaCl, 1 mM TCEP). The eluted CENP-LN complexes were concentrated and flash frozen in liquid nitrogen.

The CENP-TWSX cell pellet was resuspended in lysis buffer supplemented with 10 mM imidazole, lysed by sonication as described above and loaded onto the cOmplete His-tag purification column (Roche). The complex was eluted in 300 mM imidazole, diluted to final salt concentration of 150 mM directly and loaded onto a HiTrap Heparin HP column (Cytiva). The protein was eluted in 20 mM HEPES pH 7.8, 1 mM TCEP and 5% glycerol with a NaCl gradient from 150 mM to 600 mM. The protein was then concentrated and further purified using a Superdex 200 gel filtration column (Cytiva), Peak fractions were concentrated and flash frozen in liquid nitrogen.

To purify CENP-HIK^Head^, BL21 codon plus RIL transformants bearing pRsfDuet HIK^Head^ were grown in ZY media at 37 °C until an OD=600 nm of 0.8 was obtained, at which point expression was induced by addition of 300 μM IPTG, and cells incubated for a further 18 h overnight at 18 °C. Cells were lysed using a homogenizer (Emulsiflex, Avestin) operating at 25 kpsi, and the resulting lysate clarified by centrifugation, followed by application to StrepTactin resin (Qiagen). The resin was washed with lysis buffer, bound proteins eluted in a buffer containing 300 mM NaCl, 20 mM HEPES pH 7.8, 1 mM TCEP, 5% glycerol and the SNAP-2xStrepII tags removed overnight at 10 °C by HRC 3C protease. The CENP-HIK^Head^ complex was further purified using heparin affinity resin (Cytiva) using a linear gradient of buffer HA (20 mM Tris.HCl pH 7.5, 1 mM TCEP) against HB (1 M NaCl, 20 mM Tris.HCl pH 7.5, 1 mM TCEP), and size exclusion chromatography (Superdex S75 16/60, Cytiva; 300 mM NaCl, 20 mM HEPES pH 7.8, 1 mM TCEP). Protein fractions containing the HIK^Head^ were pooled, concentrated (Centricon, Amicon) to 15 mg/ml, flash frozen in liquid nitrogen and stored at -80 °C.

BL21 Star DE3 transformants containing GST-CENP-N^NT^ were cultured in LB at 37 °C until an OD=600 nm of 0.8 was reached. Expression was then induced overnight at 18 °C. Cells were lysed using a homogeniser (Emulsiflex, Avestin) at 25 kpsi. Clarified supernatant was loaded onto an EconoColumn (Biorad) pre-packed with GST Sepharose 4B resin (Cytiva), immobilized proteins washed with a buffer containing 300 mM NaCl, 20 mM HEPES pH 7.8, 1 mM TCEP, 5% glycerol, and eluted by addition of 5 mM reduced Glutathione. GST eluate was then purified by heparin affinity chromatography and size exclusion, exactly as described for HIK^Head^.

To purify the CENP-A octamer, fresh BL21 Star DE3 (Invitrogen) transformants harboring pRsfDuet CENP-A/H4 and pAcyc 6xHisH2A/H2B were directly inoculated into LB medium (10 colonies per litre) containing 50 μg/ml kanamycin and 25 μg/ml chloramphenicol. Cells were grown at 37 °C until reaching an OD=600nm of 0.2, at which point the temperature was reduced in two steps (24 °C until an OD=600 nm of 0.4, then 20 °C until an OD=600 nm of 0.6), and expression pursued overnight at 20 °C by induction with 300 μM IPTG.

The harvested CENP-A octamer cell pellet was lysed in a buffer of 2 M NaCl, 40 mM HEPES pH 7.8, 10 mM imidazole, 1 mM TCEP and 5 mM Benzamidine with protease inhibitors and benzonase. The octamer was eluted with a gradient of 500 mM imidazole, in lysis buffer. The eluate from Ni-NTA affinity chromatography was diluted into 500 mM NaCl with buffer of 20 mM Tris.HCl pH 7.8, 1 mM TCEP, 5 mM benzamidine. The diluted sample was further loaded onto a heparin column pre-equilibrated with the buffer of 20 mM Tris.HCl pH 7.8, 500 mM NaCl, 1 mM TCEP and 5 mM benzamidine, and the octamer was eluted by gradient to 2M NaCl in the same buffer. The complex was collected and further purified by S200 size exclusion chromatography (Cytiva), concentrated to 3 mg/ml, flash frozen in liquid nitrogen and stored at - 80 °C in the buffer of 2 M KCl, 20 mM HEPES pH 7.8, 1 mM TCEP.

The human H3 octamer was prepared by co-expression of H3, H2A, H2B and H4 in B834Rare2 cells. The harvested cell pellet was lysed in a buffer of 50 mM Tris.HCl (pH 8.0), 2 M NaCl, 1 mM EDTA and 1 mM DTT. The H3 octamer was isolated using Strep-Tactin column (Qiagen), and eluted with 4 mM desthiobiotin. The octamer was further purified by S200 size exclusion chromatography (Cytiva), concentrated to 3 mg/ml in a buffer of 10 mM Tris.HCl (pH 7.5), 2 M NaCl, 1 mM EDTA and 2 mM DTT and flash frozen in liquid nitrogen and stored at -80 °C.

### CENP-A nucleosome reconstitution

CENP-A, CENP-A mutants or H3 histone octamers were mixed with DNA fragments at 1:0.9 protein:DNA ratio with 7.8 µM final DNA concentration. The histone octamers were then wrapped by gradient dialysis from 2 M KCl to 300 mM KCl buffer with 20 mM HEPES pH 7.5, 1 mM EDTA. The KCl concentration was gradually decreased using a peristaltic pump over 16 h at room temperature. The mixture was then dialyzed for another 2 h against 20 mM HEPES pH 7.5, 300 mM KCl, 1 mM EDTA. The wrapped nucleosomes were then stored at 4 °C (after assessment).

### SEC analysis of the CCAN-CENP-A^Nuc^ complexes

To analyse assembly and stability of various CCAN and CCAN-CENP-A^Nuc^ complexes, analytical SEC experiments were performed using Superose 6 3.2/300 column (Cytiva). CCAN complexes were reconstituted at final concentration of 5 μM in CCAN gel filtration buffer (20 mM HEPES pH 7.8, 300 mM NaCl, 1 mM TCEP). The CCAN-CENP-A^Nuc^ complexes were reconstituted at 2:1 ratio (CCAN:CENP-A^Nuc^) with a final CCAN concentration of 5 μM and final nucleosome concentration of 2.5 μM. Chromatograms are plotted as absorbance at 280 nm (a.u., arbitrary unit). The eluted fractions were analyzed by 4-12% SDS-PAGE gels and stained with InstantBlue Coomassie Stain. All reconstitution assays were performed at least in duplicates.

### SEC-MALS

SEC-MALS was performed using a Wyatt MALS system. Non-crosslinked and gluteraldehyde crosslinked CCAN^ΔT^−CENP-A^Nuc^ complexes were injected onto an Superose 6 3.2/300 gel filtration column (GE Healthcare) pre-equilibrated in 20 mM HEPES (pH 7.8), 300 mM NaCl and 1 mM TCEP. The light scattering and protein concentration at each point across the peaks in the chromatograph were used to determine the absolute molecular mass from the intercept of the Debye plot using Zimm’s model as implemented in the ASTRA v.5.3.4.20 software (Wyatt Technologies). To determine inter-detector delay volumes, band-broadening constants and detector intensity normalization constants for the instrument, we used aldolase as a standard prior-to sample measurement. Data were plotted with the program Prism v.8.2.0 (GraphPad Software).

### Electrophoretic mobility shift assay (EMSA)

For CCAN-DNA EMSAs, 601 DNA was fluorescently labelled with Cy5 fluorophore at the 5’ terminus. The reaction was prepared with a constant DNA concentration of 0.5 μM and various concentrations of protein as indicated in the figures. The protein-DNA complexes were incubated in CCAN gel filtration buffer (20 mM HEPES pH 7.8, 300 mM NaCl, 1 mM TCEP) for 30 min. After addition of 5% (v/v) of glycerol, DNA and DNA-protein complexes were resolved by electrophoresis for 30 min at 88V using 0.8% TBE-agarose (w/v). The gel was visualized using Typhoon FLA 9,500 scanner (GE Healthcare) with laser set at 635 nm and Cy5 670BP30 filter. All assays were performed at least in triplicate.

For CCAN-nucleosome EMSAs, nucleosome concentration was kept constant at 2.3 μM and CCAN was added at various concentrations as indicated in the figure legends. The CCAN-nucleosome complexes were incubated in CCAN gel filtration buffer for 30 min. After addition of 5% (v/v) glycerol, nucleosome and CCAN-nucleosome complexes were resolved by gel electrophoresis for 30 min at 88V using 0.8% TBE-agarose (w/v) with added ethidium bromide. The gels were visualized using ChemiDoc XRS+ (Bio-Rad) imaging system. All assays were performed at least in triplicate.

### Quantification of the EMSA

Quantification of band intensities was performed using a Fiji scanner. Lanes for each experiment were selected and intensity estimated using a Fiji scanner. The total amount of free DNA or free nucleosome was estimated from the first lane that did not contain any additional protein for each respective repeat. The band intensity for DNA-protein or nucleosome-protein complex was then estimated, subtracting background from the first protein-free lane. To plot the fraction of DNA or nucleosome bound, the signal from the bound fraction was divided by the signal from free DNA or nucleosome fractions, respectively. The data were plotted using Prism 9 (Graphpad). Each position is the mean value from three biological replicates with one standard deviation.

### Pull-down Assays

For CENP-N^N^ binding experiments, CCAN and nucleosome were kept constant at a final concentration of 2.5 μM and 1.25 μM, respectively, in binding buffer A (200 mM NaCl, 20 mM HEPES pH 7.8, 10 mM imidazole pH 8, 1 mM TCEP, 10% glycerol, and 0.05% IGEPAL-CA630), in the combinations indicated in the figure legend, and mixed with 50 μL TALON Co^2+^ resin pre-equilibrated in binding buffer A (Takara Bio, 50% slurry) resulting in a final volume of 100 μL. Where present, CENP-N^N^ was added to a final concentration of 10 μM. Reactions were incubated with gentle rotation in falcon tubes for 30 mins at 4°C. Thereafter, 40 μL were withdrawn as an input sample, the remaining resin was washed four times with 200 μL binding buffer A, after which eluted via SDS-PAGE loading buffer, and the reactions analyzed by SDS-PAGE.

CENP-LN binding experiments were conducted as above, albeit without titration of CENP-N^N^, and using binding buffer B (150 mM NaCl, 20 mM HEPES pH 7.8, 10 mM imidazole pH 8, 1 mM TCEP, 10% glycerol, and 0.05% IGEPAL-630). Where indicated, CENP-LN was added at a concentration of 2.5 μM.

All assays were performed at least in duplicate.

### Cultivation of human cell lines and synchronization

HEK293 FlpIn-TRex cells (Invitrogen) were cultured in DMEM (Gibco) supplemented with 10% tetracycline-free FBS (PAN Biotech) at 37°C and 5% CO2. For the stable integration of eGFP-CENP-L-wt and eGFP-CENPL^cm^ into the genome, the gene was cloned into the pcDNA5-FRT-TO vector (Invitrogen) with an N-terminal eGFP-tag. HEK293 FlpIn-TREX cells were individually cotransfected with the pCDNA5-FRT-TO-CENP-L plasmids and pOG44, containing the flippase, using the HBS method. Cells were seeded the evening before transfection and the medium was exchanged the next morning. Both plasmids were mixed with 160 mM CaCl_2_ and 2×HBS buffer (final concentrations: 137 mM NaCl, 5 mM KCl, 0.7 mM Na_2_HPO_4_, 7.5 mM D-glucose and 21 mM HEPES) and added to the cells. The next steps were performed according to the Invitrogen manual. Cells were selected using 100 μg/ml Hygromycin B gold (InvivoGen). All stable cell lines were always kept under selection.

Cells were seeded in a 6-well plate in the presence of thymidine (Sigma) and doxycycline (250pg/mL, Sigma) to induce expression of the eGFP tagged transgenes for 16 h. Subsequently, cells were released by washing three times with pre-warmed medium, 4 h later the Cdk1 inhibitor RO-3306 (Santa Cruz) was added for 18 h. The cells were released again as described above for 90 min, before they were centrifuged onto poly-lysin covered cover-glasses.

### Immunofluorescence imaging

Cells on cover-glasses were pre-extracted for 2 min using PHEM + Tx (60 mM PIPES, 25 mM HEPES, 10 mM EGTA, 2 mM MgCl_2_, 2% Triton-X 100) before fixation with ice-cold methanol for 5 min at -20°C. The fixed cells were rehydrated for 10 min in PBS, permeabilized for 10 min in PBS + 0.5% Triton-X 100 and blocked in PBS, 0.5% Triton-X 100 and 2% BSA for 30 min. The following antibodies were used for staining: primary antibodies for 1 h at room temperature: CENP-A (1:500, Abcam ab13939), GFP (1:500, Abcam ab6556), secondary antibodies for 45 min at room temperature: goat anti mouse Alexa-fluor 647 and goat anti rabbit Alexa-fluor 488. The samples were also stained with 1 μg/mL Hoechst33342 (Sigma) together with the secondary antibodies. The cover-glasses were mounted using Diamond antifade (Thermo Fisher Scientific).

Images were acquired with an SP8 confocal microscope (Leica) equipped either with 405 nm, 488 nm, and 633 nm laser lines. An APO CS2 63x/1.4 oil immersion objective was used. Image resolution was set to 1024×1024 with a 3x zoom-in, bidirectional scanning, 600 Hz scan speed and 2x line average. A Z-spacing of 0.23 μm was always used. Laser power and detector gain were set for each primary antibody individually, but were kept constant between different experiments with the same antibody to ensure reproducibility.

For colocalization, images were prepared using the following workflow in ImageJ: The corresponding acquisition channels (CENP-A and eGFP) were separated and the background was subtracted from each image using the ImageJ background subtraction tool with a rolling ball radius of 5 pixels. A Gaussian blur filter was laid over the images with a sigma radius of 1. Colocalization was subsequently analyzed using the ImageJ plugin JaCoB (*57*) with the built-in object based colocalization method based on distance between geometrical centers of the identified objects. The threshold for each image was set manually to allow clear identification of kinetochores based on the CENP-A signal.

### Immunoblotting of cell lysates

Cells were seeded with a density of 0.5×106 cells/mL in a 24-well plate. Tetracycline was added and incubated for 20 h. Cells were collected by pipetting them up and down in the plate and were subsequently washed once with PBS. The cell pellet was taken up in NuPAGE LDS loading buffer (Invitrogen) and heated at 95°C for 5 min before loading onto a NuPAGE Bis-Tris 4-12% gel (Invitrogen). Immunoblotting was performed using a Mini Trans-Blot cell (BioRad) and Tris-Glycine buffer onto a nitrocellulose membrane (Amersham). Primary antibodies were incubated in 5% milk/PBS over night at 4°C and secondary antibodies in 5% Milk/PBS for 1 h at 22°C. Antibodies used were: CENP-L (Fisher PA5-114281, 1:250), GFP (Abcam ab6556, 1:1000), actin-HRP (Santa Cruz sc-47778-HRP, 1:2000), secondary anti-rabbit (Thermofisher 31462, 1:10.000).

### Crystallization, crystallography data collection and structure solution

Diffraction quality crystals of CENP-HIK^Head^ (12 mg/ml) and CENP-OPQUR^Foot^ (15 mg/ml) were obtained using the sitting drop vapor diffusion method by mixing equal volumes (200 nL) of protein and 10% PEG 8K, 100 mM imidazole pH 8.0 at 277 K or 15% PEG 2Kmme, 40 mM sodium formate, 200 mM Bis-Tris propane pH 6.9 at 18 °C, respectively. Crystals of CENP_HIK^Head^ grew after 30-60 days. CENP-OPQUR^Foot^ was treated with subtilisin in a ratio of 1:600 of enzyme per CENP-OPQUR^Foot^ complex for 30 mins at room temperature and quenched with 1 mM phenylmethylsulfonyl (PMSF) before setting up the crystallization plate. CENP-HIK^Head^ and CENP-OPQUR crystals were cryo-protected with glycerol [25% (v/v)] prior to flash freezing in liquid nitrogen. Diffraction data were collected on Diamond Light Source beamline I24 at 100 K with wavelength of 0.9795 Å, and processed with IMosflm (*58*), which allowed for precise prediction of low-resolution spots by adjusting the mosaic block size parameter from 100 to 2. Anisotropy cut-offs of diffraction limits were applied using the server STARANISO (Global Phasing). The CENP-HIK^Head^ and CENP-OPQUR^Foot^ structures were solved by molecular replacement with Phaser (*59*) using a Phyre2-generated model (*60*) and part of the cryo-EM model from this study, respectively. Interactive building was performed with COOT (*61*), refinement with REFMAC5 (*62*) and PHENIX (*63*), and validation with MolProbity (*64*). There are 0 and 0.3% Ramachandran outliers in the models of CENP-OPQUR and CENP-HIK^Head^, respectively.

### Reconstitution of the AS2-CENP-A^Nuc^-CENP-C^N^ complex for cryo-EM

CENP-C^N^ was added directly to the reconstituted AS2-CENPA^Nuc^ at 2:1 molar ratio and final nucleosome concentration of 5 µM. The sample was applied on grids directly as described below.

### Reconstitution of the CCAN^ΔT^ complex for cryo-EM

The CCAN^ΔCT^ complex was reconstituted by mixing purified CENP-OPQUR, CENP-HIKM and CENP-LN at 1:1:1 ratio at a final concentration of 5 µM. The CCAN complex was incubated for at least half an hour at room temperature. To prepare the complex for structural studies, we performed on-column glutaraldehyde (GA) cross-linking. 500 μl of 0.5% of GA was injected onto the Superose 6 Increase 10/300 column (Cytiva) with the flow rate set to 0.25 ml/min. The GA was injected for 7 ml, after which the CCAN sample was applied and eluted under an isocratic flow of 0.25 ml/min in CCAN gel filtration buffer (20 mM HEPES pH 7.8, 300 mM NaCl, 1 mM TCEP). The reaction was quenched with Tris.HCl buffer pH 8.0 at a final concentration of 50 mM straight after protein complex elution. The peak fraction of the CCAN sample was taken, concentrated and re-injected onto a Superose 6 3.2/300 column (Cytiva) coupled to a MicroÄkta (Cytiva). The resulting peak for the CCAN complex was then concentrated to 1 mg/ml and used for structural studies. For non-cross-linked CCAN^ΔT^ complex, purified CENP-OPQUR, CENP-HIKM, CENP-LN and CENP-C^N^ at 1:1:1:1 ratio were mixed together at a final concentration of 5 µM and applied on grids directly. For non-cross-linked CCAN^ΔC^ -DNA complex, the components were mixed in 1:1:1:1:1 ratio (CENP-OPQUR: CENP-LN: CENP-HIKM:CENP-TWSX:DNA) at a final concentration of 5 µM and applied on grids directly.

### Reconstitution of the CCAN^ΔT^ -CENP-A^Nuc^ complex for cryo-EM

The cross-linked CCAN^ΔT^ -CENP-A^Nuc^ complex as well as CCAN-CENP-A^Nuc^ complex was prepared identically to the cross-linked isolated CCAN^ΔCT^ complex but with addition of CENP-A nucleosome and either CENP-C^N^ protein alone or with CENP-TWSX. The protein ratios used were 1:1:1:1:2 (CENP-OPQUR: CENP-LN: CENP-HIKM:CENP-A^Nuc^:CENP-C^N^) or 1:1:1:1:1:2 (CENP-OPQUR: CENP-LN: CENP-HIKM:CENP-TWSX:CENP-A^Nuc^:CENP-C^N^), to a final nominal concentration of 5 µM.

### Cryo-EM grid preparation and data acquisition

3 µl of prepared complexes at a concentration of approximately 1 mg/ml was applied to a glow-discharged Quantifoil 300 mesh copper R1.2/1.3 grids (Quantifoil Micro Tools). The grids were then flash frozen in liquid ethane using a Thermo Fisher Scientific Vitrobot IV (0-0.5 s waiting time, 2 s blotting time, -7 blotting force). Cryo-EM images were collected on Thermo Fisher Scientific Titan Krios microscope operating at 300 keV using a Gatan K3 camera. A magnification of 105 k was used for the apo-CCAN^ΔCT^, and CCAN-CENP-A^Nuc^-AS1 samples, yielding pixel sizes of 0.86 Å/pixel and 0.831 Å/pixel, respectively. Additional data were collected for CCAN-CENP-A^Nuc^-AS1 at a magnification of 81 k, yielding nominal pixel sizes of 0.93 Å. The images were recorded at a dose rate of 25 e^-^/px/s with 2 s exposure and 60 frames. For AS2-CENPA^Nuc^-CENP-C^N^, CCAN^ΔC^ -DNA complex and CCAN-CENP-A^Nuc^ complex, a magnification of 105 k was used yielding pixel sizes of 0.853 Å/pixel. The images were recorded at a dose rate of 16 e^-^/px/s with 2.1 s exposure and 40 frames. The Thermo Fisher Scientific automated data-collection program EPU was used for data collection, with AFIS. Defocus values ranged from -1.6 to -3.0 µm, at an interval of 0.2 µm.

### Cryo-EM data processing

For the CCAN^ΔCT^ -DNA, AS2-CENP-A^Nuc^-CENP-C^N^, CCAN^ΔC^-DNA, AS1-CENP-A^Nuc^-CENP-C, and CCAN^ΔT^-CENP-A^Nuc^-AS2 reconstructions, micrograph movie frames were aligned with MotionCor2 (*65*), and contrast-transfer function (CTF) estimation was performed by CTFFIND (*66*), as integrated into RELION3.1 (*67*). Particle picking was performed with SPHIRE-crYOLO (*68*). For the apo-CCAN and CCAN^ΔT^-CENP-A^Nuc^-AS1 reconstructions, the same processing steps were instead carried out with Warp (*69*).

For apo-CCAN, reference-free 2D classification in RELION yielded classes with clear secondary structure which were then exported to cryoSPARC (*70*) for *ab initio* model generation, using a model number of 2. The resulting map was then used in 3D classification in RELION (*67*). Particle classes bearing the most homogeneous density were then subjected to 3D refinement, followed by iterative CTF parameter refinement and Bayesian polishing. Consensus refinement thus yielded a volume with an overall resolution of 3.2 Å. Multibody refinement was then applied using partially overlapping masks around the CENP-OPQURN, and CENP-HIKMLN lobes, yielding resolutions of 3.9 Å, and 3.2 Å respectively. To further improve the continuity of the CENP-OPQURN density, particle subtraction was applied to remove the CENP-HIKML signal, and particles were exported to cryoSPARC, where they were further heterogeneously classified against a true and two noisy decoy volumes. Resulting particles were then reimported into RELION (*67*) and refined, yielding a 3.6 Å volume. To derive volumes with more continuous CENP-HIK^Head^ density, the CENP-OPQURN signal was subtracted, and resulting particles classified in 3D without alignment, using a T value of 8. Particle classes corresponding to the raised and lowered states of CENP-HIK^Head^ were then refined, producing reconstructions with resolutions of 4.87 Å and 4.11 Å. Reconstruction of the full raised and lowered CCAN volumes was achieved by reverting the subtraction of refined particle sets, and a final round of consensus 3D refinement.

For CCAN^ΔT^ -CENP-A^Nuc^, rigorous 2D classification, in RELION, was essential to identify complexes containing all constituents. Classes with clear features corresponding to CCAN-DNA and CENP-A^Nuc^ were exported to cryoSPARC (*70*) for *ab initio* model generation, as for apo-CCAN. The particle set arising from 2D classification was then classified iteratively in 3D against the resulting models, in RELION. 3D consensus refinement of the best-defined classes produced an initial reconstruction with an overall resolution of 10 Å. Signal subtraction of the CENP-A^Nuc^ component of these particles, followed by 3D classification with alignment, and 3D consensus refinement, yielded a reconstruction of the CCAN-DNA subdomain with an overall resolution of 8.7 Å, enabling assignment of the register of DNA emanating from the CENP-A nucleosome. To further improve the resolution of the CCAN-CENP-A^Nuc^ map, particles originating from two data collection sessions were scaled to the same pixel size (*71*), 1.662 Å, and exported to cryoSPARC. Two rounds of further heterogeneous refinement against two noise decoys and a single true map were performed, followed by homogeneous and non-uniform refinement (*72*), yielding a CCAN^ΔT^ -CENP-A^Nuc^ volume with an overall resolution of 8.72 Å. All resolution values were estimated according to the Gold-Standard Fourier Shell Correlation (GS-FSC) = 0.143 criterion. Volume post-processing and local resolution estimation were performed in RELION. Identical procedures were follwoed for the CCAN -CENP-A^Nuc^, resulting in a density volume with an overall resolution of 12 Å.

For CCAN^ΔT^-DNA and CENP-A^Nuc^-CENP-C^N^ complexes, data were initially subjected to 2D classification in cryoSPARC. Classes corresponding to CCAN and CENP-A nucleosomes were picked and *ab initio* models generated. *Ab initio* models were further refined individually to obtain better-defined cryo-EM densities. Heterogeneous refinement in cryoSPARC with the refined CENP-A^Nuc^-CENP-C^N^ volume, CCAN-DNA volume and four decoy noise volumes was performed against the entire set of automatically picked particles. Particles corresponding to CENP-A^Nuc^-CENP-C^N^ and CCAN-DNA were selected and kept separate in the subsequent data processing workflow. Another round of heterogeneous refinement was performed using either CENP-A^Nuc^-CENP-C^N^ volume or CCAN-DNA volume and five dummy noise volumes for each particle set. For CENP-A^Nuc^-CENP-C^N^, particles were then exported back to RELION 3.1 and two successive rounds of CTF refinement and Bayesian polishing were performed, and a final, polished particle dataset was used a final round of 3D refinement in RELION, yielding a final resolution of 2.7 Å (GS-FSC). For CCAN-DNA, the particle set was further cleaned by 2D classification, and the best 2D class averages with well-resolved secondary structure were used for two successive rounds of non-uniform refinement in cryoSPARC, yielding a final resolution of 4.5 Å.

For CCAN^ΔC^-DNA, 2D classification was performed in cryoSPARC. The best classes corresponding to CCAN^ΔC^-DNA were selected and initial model generated *ab initio. Ab initio* models were further refined individually to obtain better-defined cryo-EM densities. Two rounds of heterogeneous refinement were performed in cryoSPARC with refined CCAN^ΔC^-DNA volume and five decoy noise classes against the entire dataset of automatically picked particles. The best particles were subsequently refined in RELION 3.1 with two successive rounds of CTF refinement and Bayesian polishing. The final, polished dataset was used in a final round of 3D refinement in RELION to obtain a final resolution of 2.83 Å.

For AS2-CENP-A^Nuc^-CENP-C^N^ 2D classification was performed in cryoSPARC. The best classes were selected and initial model generated ab initio. *Ab initio* models were further refined individually to obtain better-defined cryo-EM densities. One round of heterogeneous refinement were performed in cryoSPARC with refined AS2-CENP-A^Nuc^-CENP-C^N^ volume and five decoy noise classes against the entire dataset of automatically picked particles. 2D classification was used to select the best resolved classes. The best particles were refined using homogeneous and non-uniform refinement in cryoSPARC to obtain a final resolution of 2.44 Å.

CryoSPARC implementation of 3D variability analysis (3DVA) was used to understand conformational heterogeneity of the AS2-CENP-A^Nuc^-CENP-C^N^ complex. 3 modes were analysed with 250k particles as initial input and resolution limit of 7 Å. The results were visualized in Chimera by splitting the first mode of 3DVA into 20 clusters (Supplementary movie 1).

### Cryo-EM model building and refinement

#### Apo-CCAN

Homology models for CENP-H, CEHP-I, CENP-K and CENP-L were generated by one-to-one threading in Phyre2 (*60*), based on *S. cerevisiae* CCAN (*23*) (PDB 6QLE). Crystal structures were available for CENP-N^NT^ (*8*) (PDB 6EQT) and CENP-M (*24*) (PDB 4P0T). These models were docked into the corresponding map density by rigid body fitting in Chimera and COOT. CENP-O, CENP-P, CENP-Q, CENP-U, CENP-R were built *de novo*. Models encompassing the entire CCAN^ΔCT^ were manually rebuilt, extended, and corrected in COOT (*61*), followed by real-space refinement in PHENIX (*63*) against a composite map. The crystal structure of CENP-OPQUR aided the building of the CENP-OPQUR module in the cryo-EM maps. We further compared and modified our models according to AlphaFold2 (*36, 73*) predictions, which were released in the course of this study. The AlphaFold2 predictions particularly aided the building of the C-terminal regions of CENP-Q and CENP-U and CENP-R.

#### CCAN-DNA

Models for CCAN^ΔCT^ and ideal B-form DNA were rigid-body fitted into the corresponding volumes in Chimera, followed by manual correction in COOT (*61*) and real-space refinement in PHENIX (*63*).

#### CENP-A^Nuc^-CENP-C

PDB ID 6MUO model (*11*) was used as the initial model. The model was fitted into the cryo-EM map in ChimeraX (*74*), manually corrected in COOT (*61*) and refined in PHENIX (*63*). The DNA sequence was manually re-build in COOT (*61*).

#### CCAN-CENP-A^Nuc^

Higher resolution density maps of CENP-A^Nuc^, isolated CCAN^ΔT^-DNA, and the CCAN^ΔT^-DNA volume resulting from focused refinement were fitted into the 10 Å consensus volume for the complete CCAN-CENP-A^Nuc^ complex, and resampled on a common origin. Models for CCAN^ΔCT^, and CENP-A^Nuc^-CENP-C were rigid-body fitted into the corresponding map density regions. The phase of the DNA gyre emerging from the nucleosome could be readily discerned, and was modelled through manual building and real-space correction in COOT (*61*), invoking chain self-restraints to prevent distortion of proper geometry at low resolution.

### AlphaFold2 Predictions

AlphaFold2 (*36*) predictions were run using versions of the program installed locally and on ColabFold with the MMseqs2 MSA option. For the CENP-C^PEST^ – CENP-LN prediction, input sequences were CENP-C (residues 201-400), CENP-N (221 to C-term), CENP-L (all residues); and for CENP-C^PEST^ – CENP-HIKM, input sequences were: CENP-C (residues 101-350), CENP-H (residues 1-190), CENP-I (residues 341 to C-term), CENP-K (residues 1-160), CENP-M (all residues). For CENP-OPQUR prediction, input sequences were: CENP-O and CENP-P (all residues), CENP-Q (residues 193 to C-term), CENP-U (residues 351 to C-term), CENP-R (all residues).

---

**Fig. S1.**
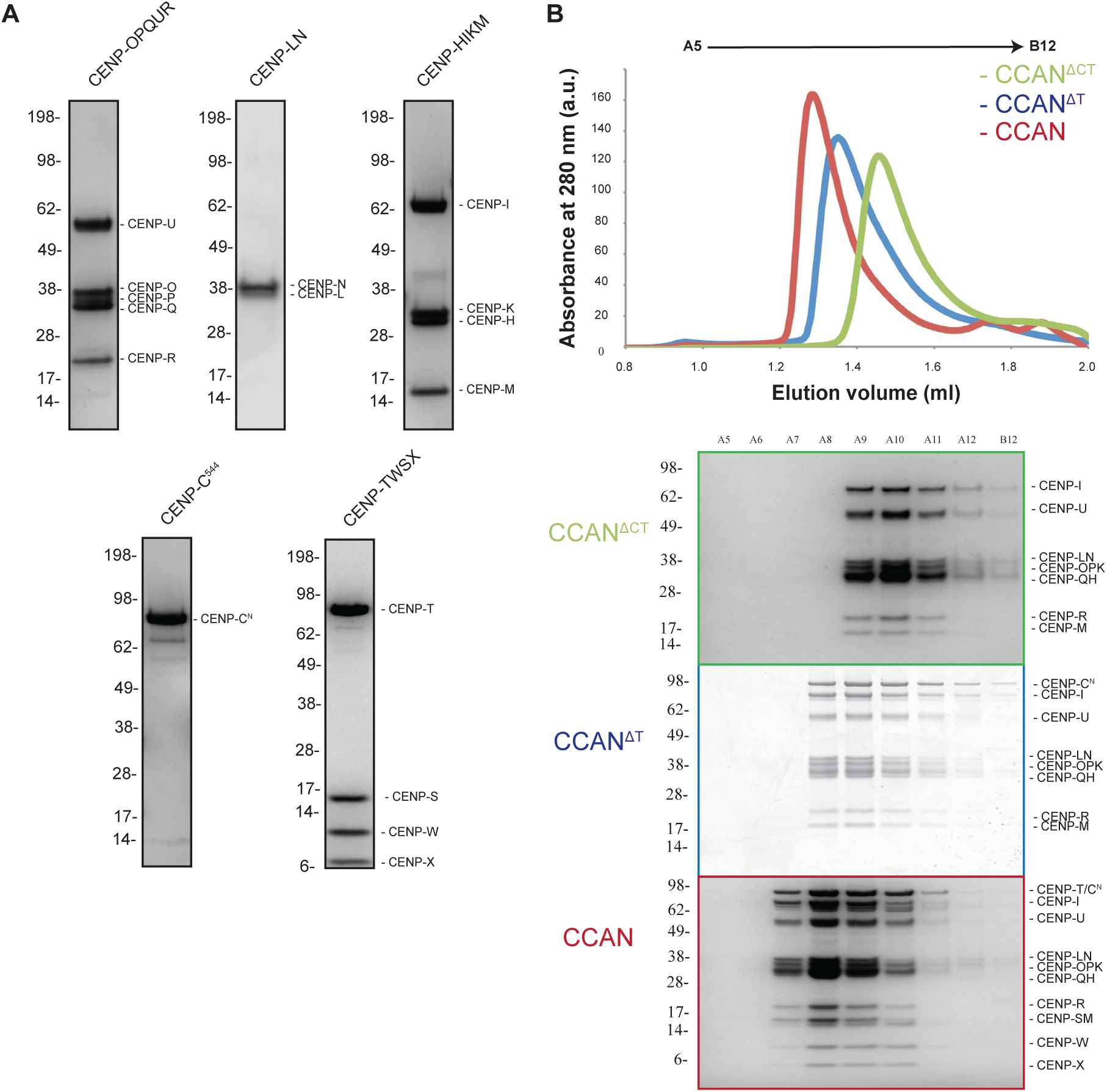
Reconstitution of the human CCAN complexes. (**A**) Coomassie-blue-stained SDS-PAGE gels of the purified CCAN modules used in this study. (**B**) Upper panel: Size-exclusion chromatography profiles (Superose 6 Increase 3.2/300 (Cytiva)) of the reconstituted CCAN, CCAN^ΔT^ and CCAN^ΔCT^ complexes showing single high molecular weight peaks for each complex. Lower panel: Coomassie-blue-stained SDS-PAGE gels of the reconstituted CCAN complexes showing that peak fractions contain all of the CCAN modules used in the reconstitution.

**Fig. S2.**
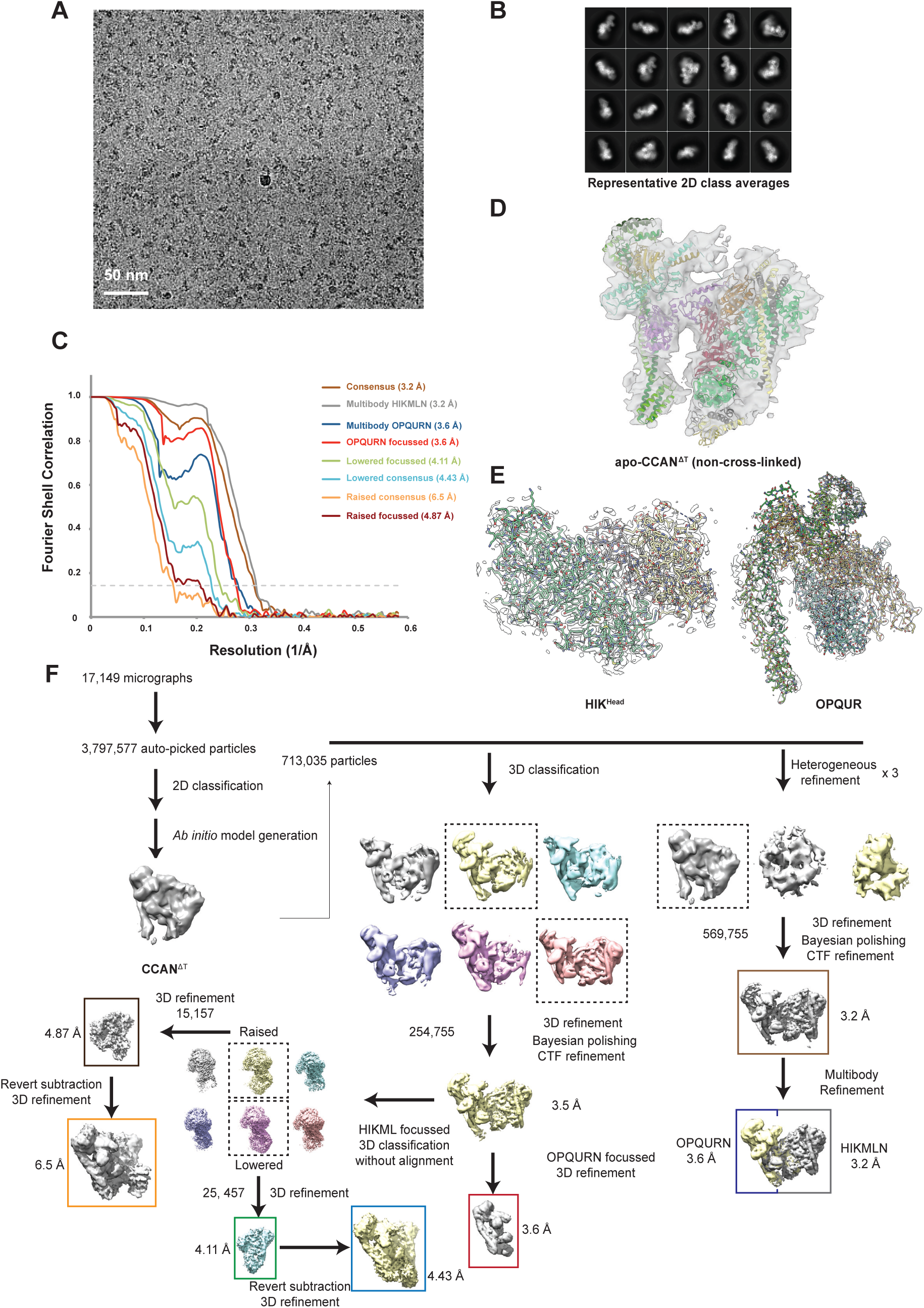
Cryo-EM data of the human apo-CCAN complexes. (**A**) A typical cryo-electron micrograph of apo-CCAN^ΔCT^, representative of 17,149 micrographs. (**B**) Galleries of 2D class averages of apo-CCAN^ΔCT^, representative of 100 2D classes. (**C**) FSC curves shown for the cryo-EM reconstructions of apo-CCAN^ΔCT^. FSC curves are defined in panel F. (**D**) Cryo-EM reconstruction of uncross-linked apo-CCAN^ΔT^ with fitted atomic model of CCAN^ΔCT^. (**E**) Crystal structures of CENP-HIK^Head^ (left) and CENP-OPQUR (right) with associated 2Fo-Fc electron density map. (**F**) Workflow for cryo-EM data processing of apo-CCAN.

**Fig. S3.**
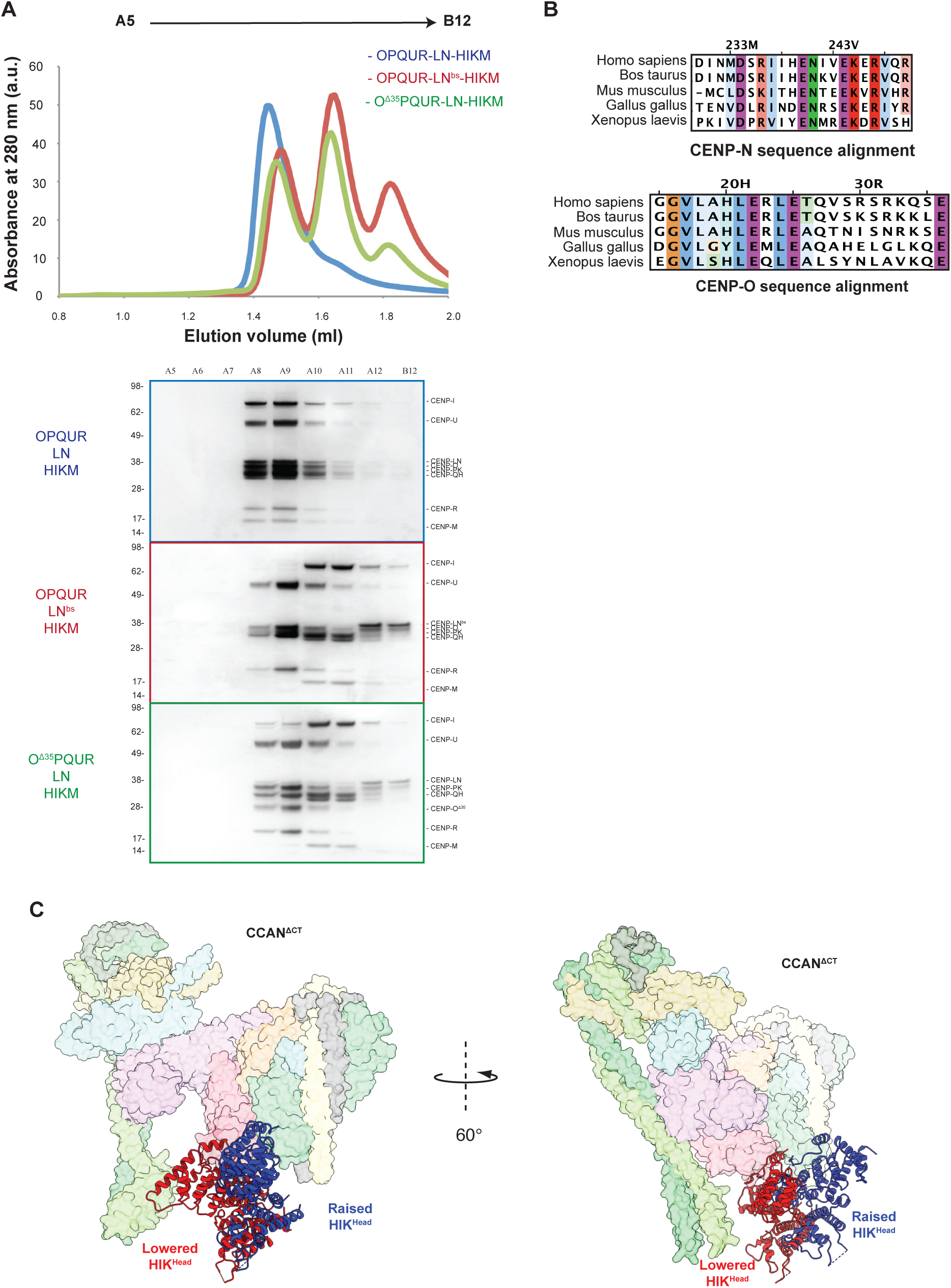
Testing the assembly principles of the CCAN^ΔCT^. (**A**) Comparative SEC profiles (Superose 6 increase 3.2/300 (Cytiva)) of either fully assembled or mutated CCAN^ΔCT^, composed of CENP-OPQUR, CENP-LN and CENP-HIKM modules. The two mutants (i) CENP-N β-strand mutant (CENP-N^bs^) where 236-240 amino acids were replaced with alanines, and (ii) CENP-O mutant where the N-terminal 35 amino acids were deleted (CENP-O ^Δ35^ mutant) show defects in apo-CCAN^ΔCT^ assembly, suggested by the disassembly of the apo-CCAN^ΔCT^ into three stable modules. Lower panel: Coomassie-blue-stained SDS-PAGE gels for each experiment confirmed the composition of each peak. (**B**) Multiple sequence alignments (MUSCLE) of the β-strand of CENP-N, and N-terminus of CENP-O shows a high degree of sequence conservation in these regions of CCAN. (**C**) Two conformational states (raised and lowered) of the CENP-HIK^Head^ sub-module.

**Fig. S4.**
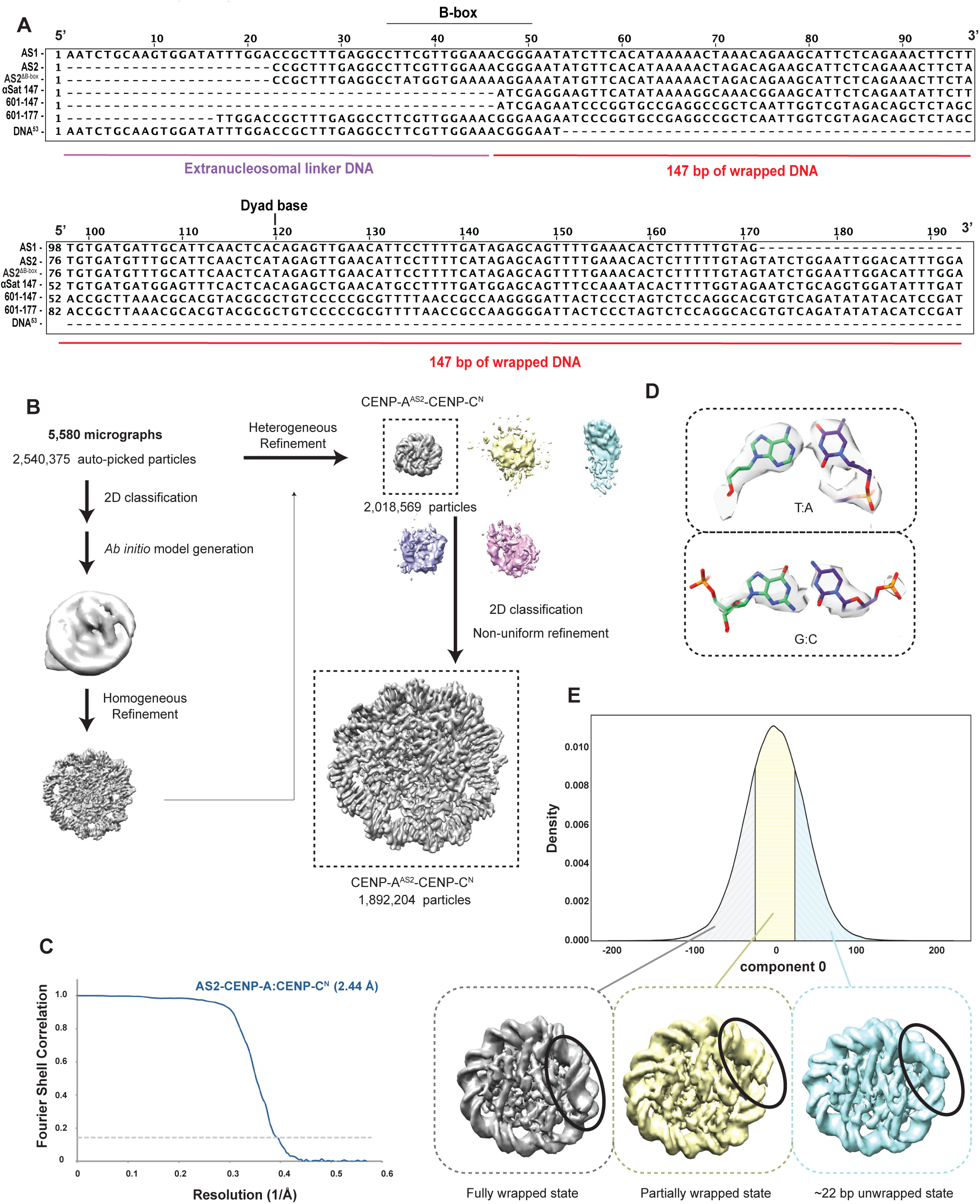
Reconstitution of the CCAN^ΔT^-CENP-A^Nuc^ complex. (**A**) MUSCLE sequence alignment of all the nucleosome reconstitution DNA constructs that were used in this study to reconstitute CCAN-CENP-A^Nuc^ complexes. The 5’ and 3’ ends of the DNA referred to in the text are indicated. Numbering scheme is based on the *α*-satellite repeat defined by (*26*). (**B**) Workflow for cryo-EM structure determination of the AS2-CENP-A^Nuc^-CENP-C^N^ complex. (**C**) FSC curve for the AS2-CENP-A^Nuc^-CENP-C^N^ complex. (**D**) Representative density for the nucleotide base pairs wrapped around the CENP-A nucleosome. (**E**) The relative probabilities of observing the nucleosome in wrapped (grey, left), intermediate (yellow, middle) and unwrapped (cyan, right) states by kernel density estimation of the eigenvalues for the first eigenvector in a 3D variability analysis experiment in cryoSPARC.

**Fig. S5.**
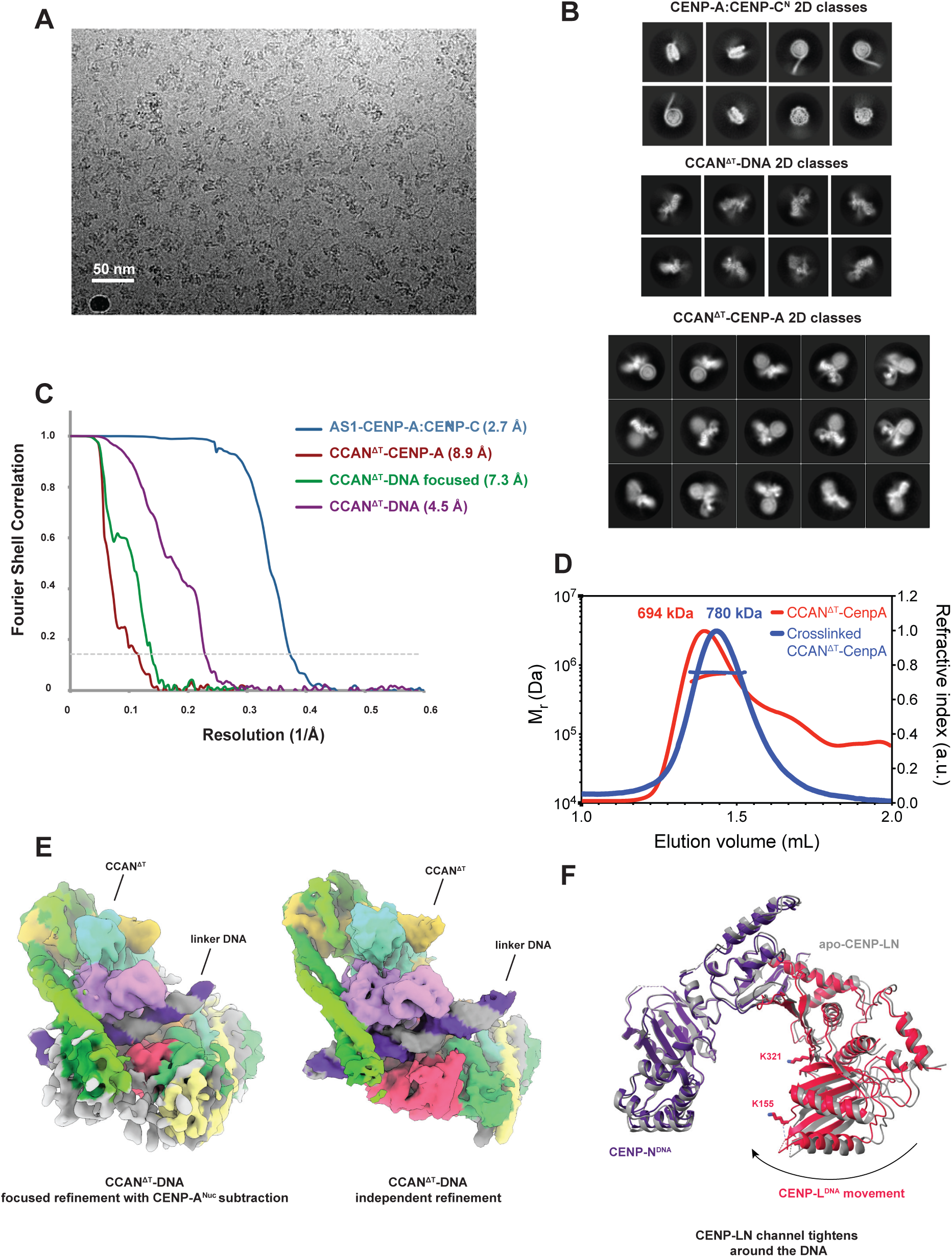
Cryo-EM data of the human CCAN^ΔT^ complexes with CENP-A^Nuc^, CENP-C^N^ and DNA (AS1 DNA sequence). (**A**) A typical cryo-electron micrograph of CCAN^ΔT^-CENP-A^Nuc^ AS1 complex, representative of 38,345 micrographs. (**B**) Galleries of 2D class averages of CENP-A^Nuc^-CENP-C^N^ (top panel), CCAN^ΔC^-DNA (middle panel) and CCAN^ΔCT^-CENP-A^Nuc^ (bottom panel), representative of 100 2D classes. (**C**) FSC curves shown for the cryo-EM reconstructions of CCAN^ΔT^ complex. FSC curves are defined in **Figure S6**. (**D**) Representative SEC-MALS data for non-cross-linked as well as cross-linked CCAN^ΔT^-CENP-A^Nuc^ complex. The predicted mass of the CCAN^ΔT^-CENP-A^Nuc^ complex is 710 kDa, the predicted mass of CCAN^ΔT^-CENP-A^Nuc^ with two copies of CENP-C^N^ is 796 kDa. (**E**) Cryo-EM maps comparing CCAN^ΔT^-CENP-A^Nuc^ and CCAN^ΔCT^-DNA showing the similar mode of linker DNA interaction in the two complexes. (**F**) Superimposition of the CENP-LN module in apo-CCAN (grey) onto the CCAN-DNA complex (indigo and red) shows that the CENP-LN channel contracts when bound to DNA.

**Fig. S6.**
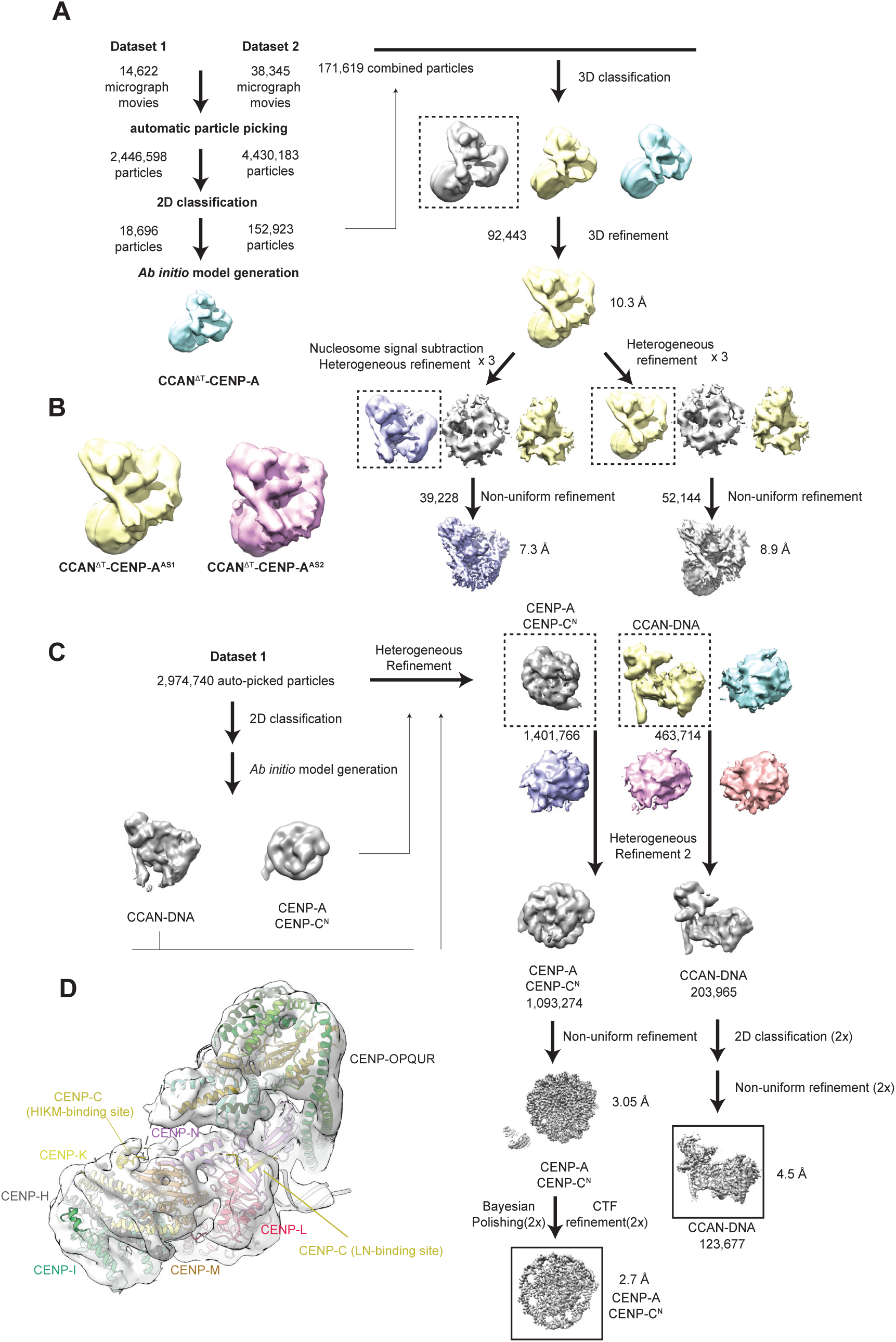
Workflow for cryo-EM reconstructions of CENP-A^Nuc^-CENP-C^N^, CCAN^ΔT^-DNA and CCAN^ΔT^-CENP-A^Nuc^ complexes. (**A**) Workflow for cryo-EM reconstruction of CCAN^ΔT^-CENP-A^Nuc^ for the AS1 DNA sequence (CCAN^ΔT^-CENP-A^AS1^). An identical work-flow was applied for CCAN^ΔT^-CENP-A^AS2^. (**B**) The structures of CCAN^ΔT^-CENP-A^Nuc^ reconstituted with the two *α*-satellite repeat DNA sequences AS1 and AS2 (defined in **fig. S4A**) are essentially identical. For the CCAN^ΔT^-CENP-A^AS2^ complex, the dataset comprised 33,097 micrographs and 53,292 particles were used in the final reconstruction. (**C**) Workflow for cryo-EM reconstruction of CCAN^ΔT^-DNA and CENP-A^Nuc^-CENP-C^N^. (**D**) AlphaFold2 generated models of CENP-C^PEST^ interactions with the CENP-LN and CENP-HIKM modules readily fitted into the unassigned EM density in the CCAN^ΔT^-DNA complex.

**Fig. S7.**
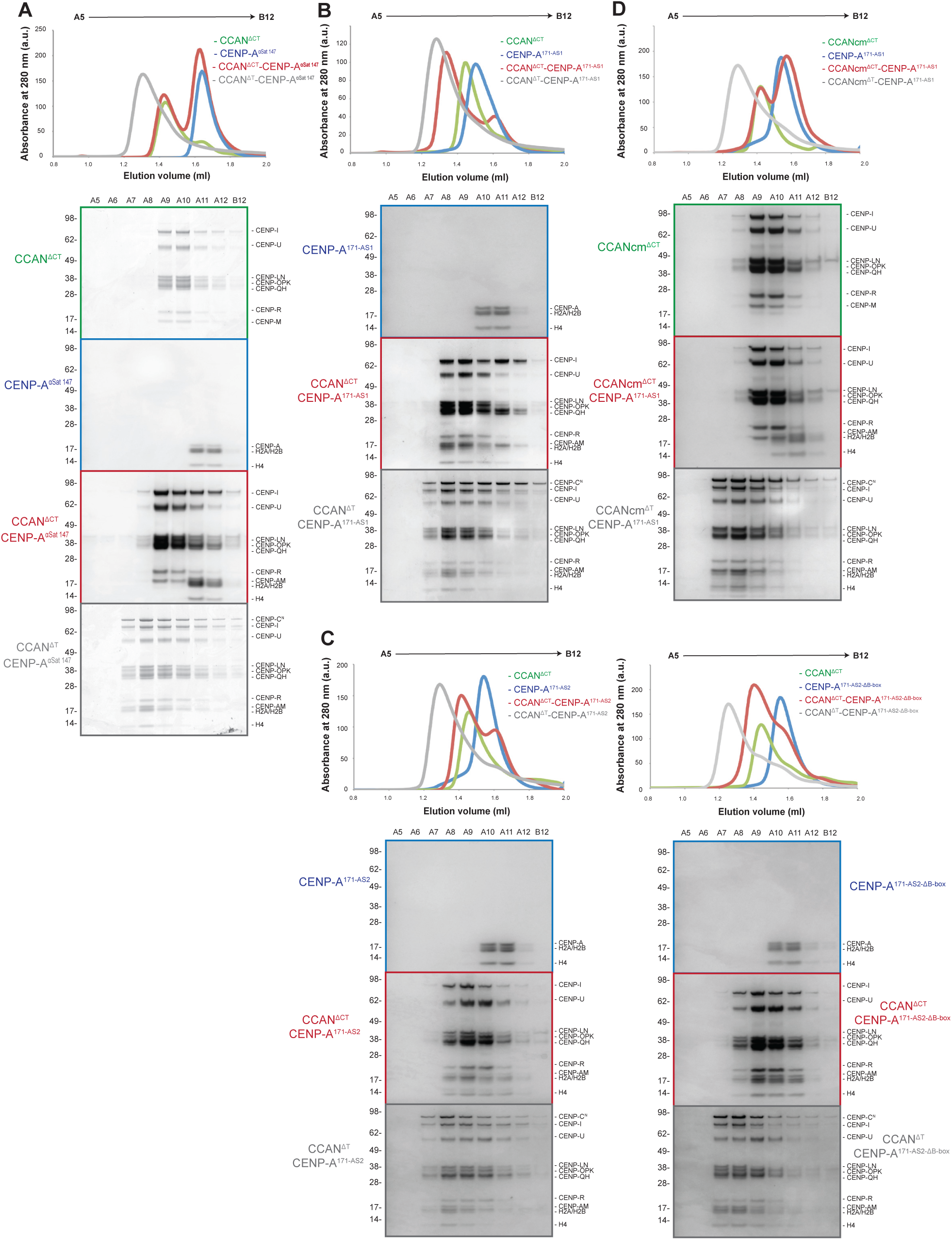
Reconstitution of the CCAN^ΔCT^ and CCAN^ΔT^ with CENP-A^Nuc^ wrapped using different DNA substrates. (**A-D**) Comparative SEC profiles (Superose 6 increase 3.2/300 (Cytiva)) of either CCAN^ΔCT^ or CCAN^ΔT^ with CENP-A^Nuc^ and their corresponding Coomassie-blue-stained SDS-PAGE gels. All reconstitutions were performed at least in two independent duplicates. (**A**) CCAN^ΔCT^ does not bind to CENP-A^Nuc^ reconstituted with αSat147 sequence as seen by elution of two separate complexes (CCAN^ΔCT^-CENP-A*^α^*^Sat-147^). Addition of CENP-C^N^ allowed formation of the CCAN^ΔT^-CENP-A^Nuc^ complex with the *α*Sat147 DNA sequence (CCAN^ΔT^-CENP-A*^α^*^Sat-147^). (**B**) CCAN^ΔCT^ readily bound to CENP-A^Nuc^ reconstituted with AS1 DNA sequence (αSat171 based). The binding was further augmented by addition of CENP-C^N^. (**C, D)** CCAN^ΔCT^ bound to CENP-A^Nuc^ reconstituted with the AS2 DNA sequence (αSat171 based) and AS2 DNA sequence which lacks the B-box (AS2**^Δ^**^B-box^). The binding was further strengthened by addition of CENP-C^N^. (**D**) CCAN^ΔCT^ charge mutant (CCANcm^ΔCT^), in which the positive patch on the CENP-L backside proximal to the nucleosomal DNA gyre (R46, R47, K48) as well as two positively-charged residues lining the central DNA-binding groove (K155, K321), were mutated to invert the charge, assembled into CCANcm^ΔCT^ yet failed to bind to the CENP-A^Nuc^ reconstituted with the AS1 DNA sequence. Addition of CENP-C restored binding.

**Fig. S8.**
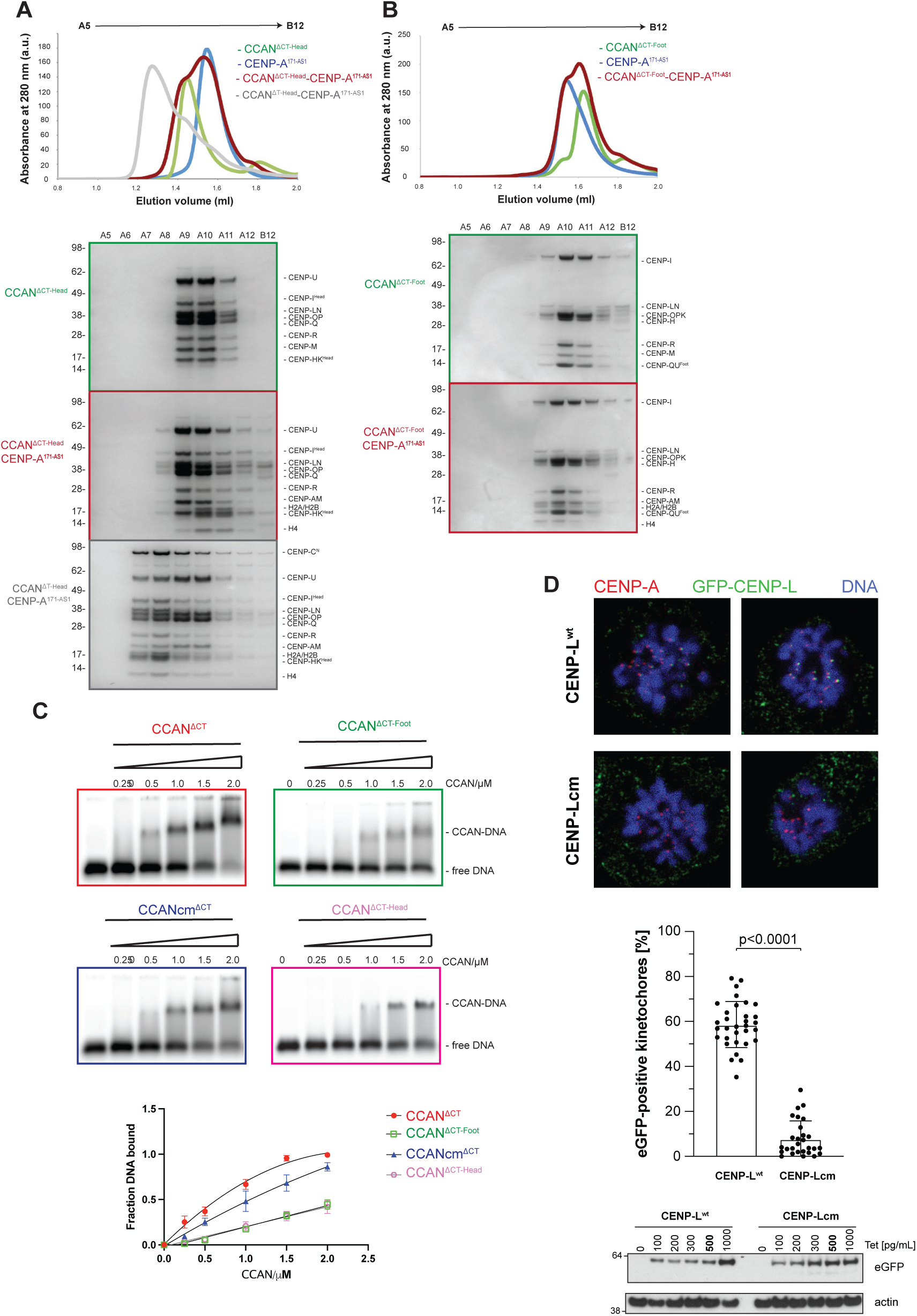
Multiple domains of the CCAN are necessary for stable binding to the CENP-A^Nuc^. (**A, B**) Comparative SEC profiles (Superose 6 increase 3.2/300 (Cytiva)) of CCAN^ΔCT^ mutants where either HIK^Head^ group or OPQUR^Foot^ were deleted and their corresponding Coomassie-blue-stained SDS-PAGE gels below. Deletion of either HIK^Head^ group or OPQUR^Foot^ abolished binding of CCAN^ΔCT^ to CENP-A^Nuc^. All reconstitutions were done at least in two independent duplicates. (**C**) EMSA assays between either wild-type and mutant CCAN^ΔCT^ and free 601 Widom DNA (147 bp) labelled with Cy5 dye. The fraction of bound DNA was quantified across three independent replicates as described in the Methods and plotted as a mean value with one standard deviation. (**D**) Top panel: mitotic HEK293 cells expressing either eGFP-CENP-Lwt or eGFP-CENP-Lcm were fixed and stained using the indicated antibodies. Exemplary single Z-stacks from two individual cells each are shown to illustrate co-localization of the eGFP tagged constructs with kinetochores marked by CENP-A. The co-localization of the eGFP tagged constructs with kinetochores was analysed. N = 32 cells for CENP-L^wt^ and 28 cells for CENP-Lcm. P-value was calculated using a Mann-Whitney U-test. Bottom panel: Whole cell lysate was prepared from HEK293 cells expressing either eGFP-CENP-L^wt^ or eGFP-CENP-Lcm. Immunoblotting was performed against the indicated proteins. A tetracycline (Tet) concentration of 500pg/mL was used for all experiments with these cell lines.

**Fig. S9.**
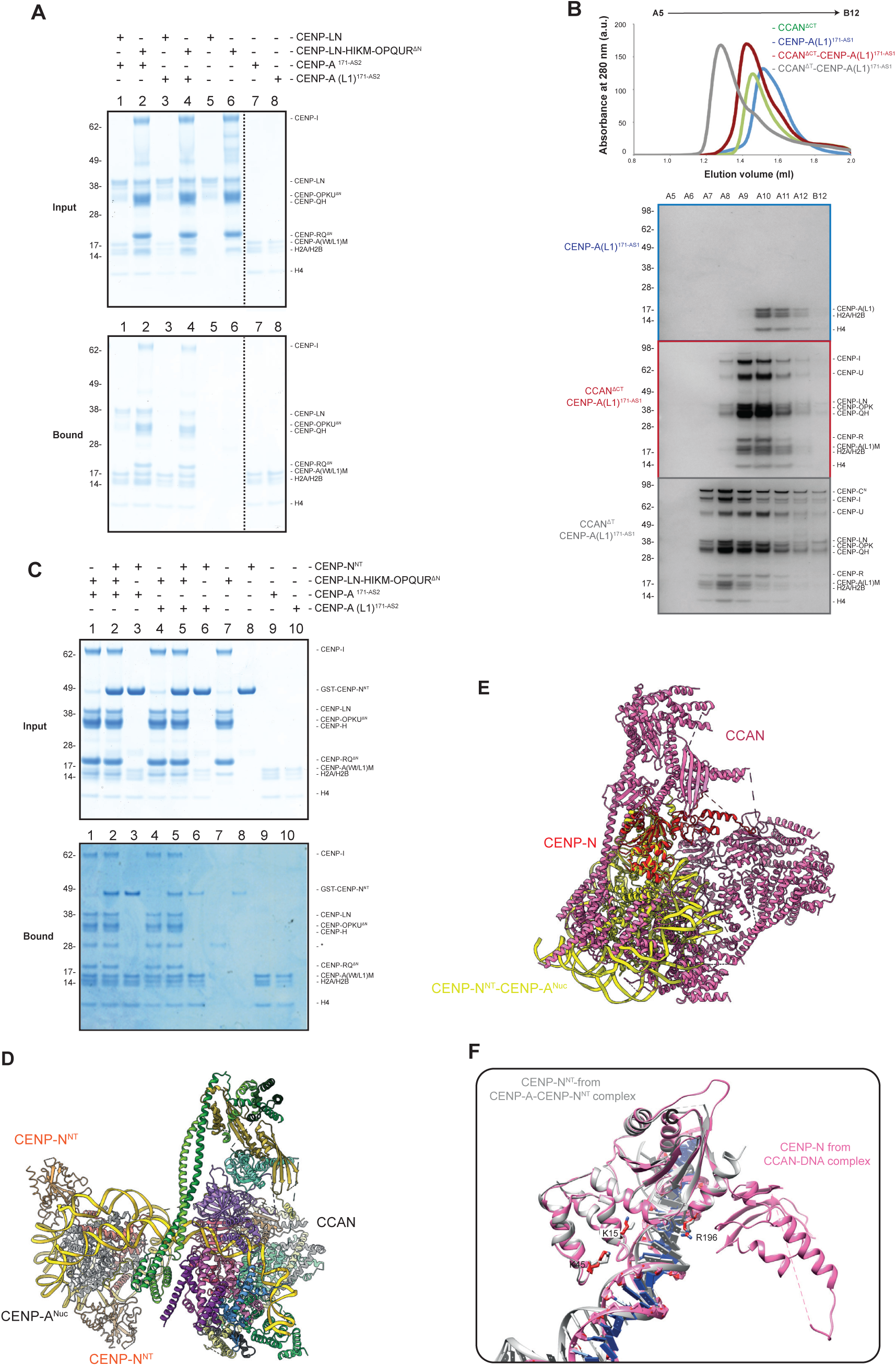
Assembly of CCAN on the CENP-A nucleosome is not dependent on the L1-loop. (**A**) Pulldown assays of CENP-LN and CENP-LN-HIKM-OPQUR*^Δ^*^N^ against CENP-A^Nuc^ wild-type or L1 loop mutant, in which the L1-loop RG motif was mutated to AA, labeled CENP-A (L1), at 150 mM NaCl. A dashed line indicates the boundary of cropped gel images. While wild-type CENP-A^Nuc^ retains CENP-LN (lanes 1), binding is entirely abolished by the L1-loop mutation (lane 2). In contrast, CENP-LN-HIKM-OPQUR*^Δ^*^N^ binds equally well to either nucleosome, confirming that CCAN assembly onto the nucleosome does not require the L1-loop: lanes 2 (wild-type CENP-A), and 4 (L1-loop mutant CENP-A). A gently truncated version of CENP-QU N-termini (CENP-OPQUR*^Δ^*^N^, see Methods) had to be used in these assays as full-length CENP-QU is insoluble under this salt condition. (**B**) Comparative SEC profiles of CCAN^ΔCT^ and CCAN^ΔC^ reconstituted with a CENP-A^Nuc^ (L1). CENP-A(L1)^Nuc^ was reconstituted with the AS1 DNA sequence. SEC as well as Coomassie-blue-stained SDS-PAGE gels indicated that both CCAN^ΔCT^ and CCAN^ΔT^ formed stable complexes with CENP-A(L1)^Nuc^. (**C**) Pulldown assays of GST-CENP-N^NT^ and CENP-LN-HIKM-OPQUR*^Δ^*^N^ against wild-type and L1 loop mutant variants of CENP-A^Nuc^, at 200 mM NaCl. CENP-N^NT^ is dependent on the L1 loop for nucleosome binding: lanes 3 (wild-type CENP-A), 6 (L1-loop mutant CENP-A), and lanes 8 (GST-CENP-N resin binding control). Addition of a four-fold molar excess of GST-CENP-N^NT^ over CENP-LN-HIKM-OPQUR*^Δ^*^N^ does not impede CCAN retention by CENP-A nucleosomes, however binding of the GST-CENP-N^NT^ but not the CENP-LN-HIKM-OPQUR*^Δ^*^N^ is still dependent on the L1-loop: lanes 1/2 (binding of CENP-LN-HIKM-OPQUR*^Δ^*^N^ to wild-type CENP-A^Nuc^ in the absence and presence of GST-CENP-N^NT^, respectively), and lanes 4/5 (same experiment, albeit with L1-loop mutant CENP-A^Nuc^). A contaminant band is indicated with an asterisk. (**D**) Modelling shows that CCAN and CENP-N^NT^ can simultaneously bind to a CENP-A^Nuc^-171 bp module with CCAN interaction with extranucleosomal DNA and CENP-N^NT^ interactions with the CENP-A^Nuc^ L1 loop. (**E**) Superimposing CENP-N^NT^ of the CENP-N^NT^-CENP-A^Nuc^ complex (*8–11*), onto CENP-N^NT^ of CCAN in the context of CCAN-CENP-A^Nuc^ (this study) shows that CENP-L, CENP-HIKM and CENP-QU would clash extensively with CENP-A^Nuc^ of the CENP-N^NT^-CENP-A^Nuc^. **(F**) Comparative figure showing that the same basic residues of CENP-N in either free CENP-N^NT^ in the CENP-N^NT^-CENP-A^Nuc^ complex, or in the context of CCAN in the CCAN-CENP-A^Nuc^ complex (this study) contact the DNA gyre of CENP-A^Nuc^, although at different SHLs.

**Fig. S10.**
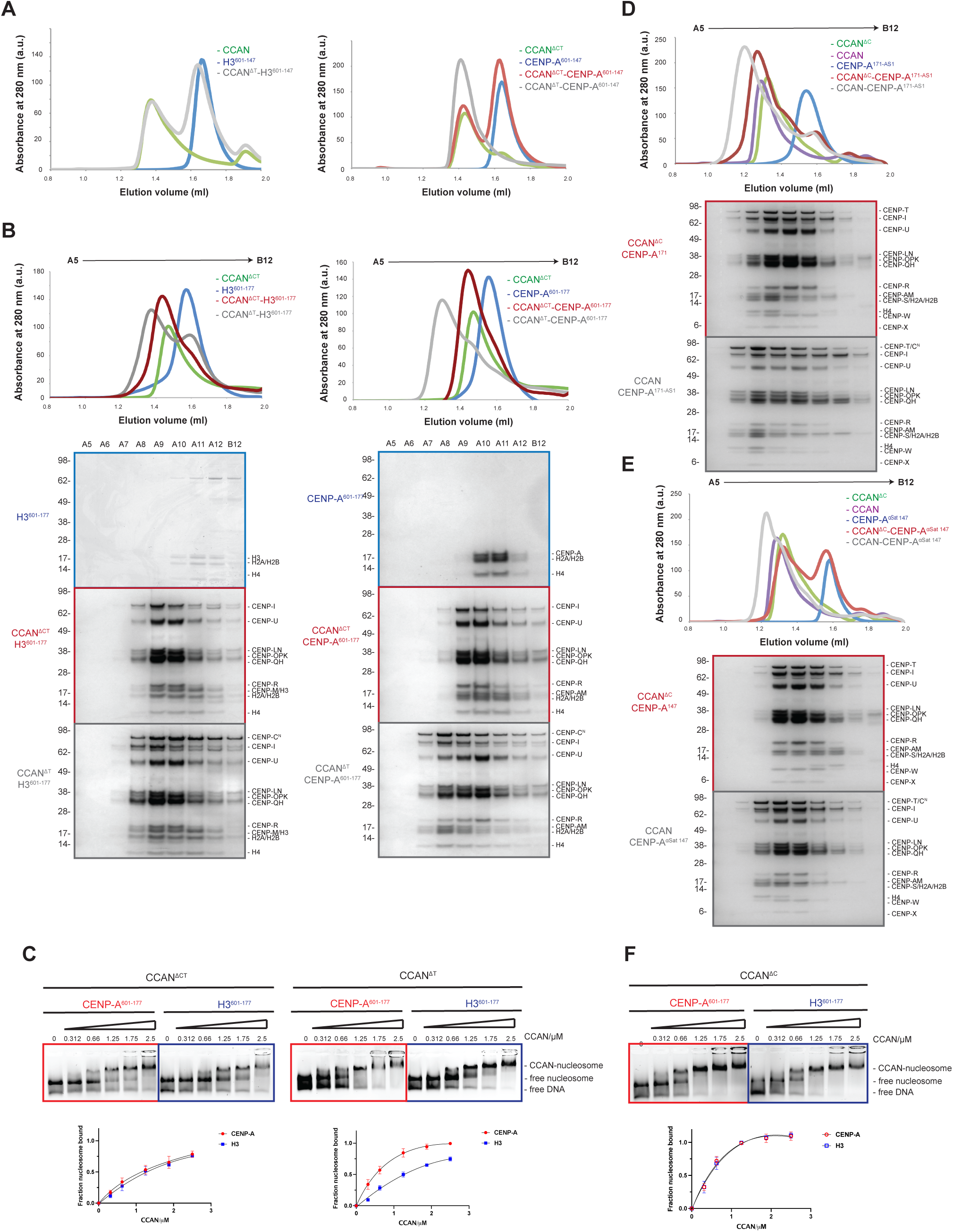
The selectivity between CCAN and CENP-A^Nuc^ is conferred by CENP-C^N^ and reconstitution of the complete CCAN-CENP-A^Nuc^ complex. (**A**) Comparative SEC profiles (Superose 6 increase 3.2/300 (Cytiva)) of CCAN binding to either H3^Nuc^ or CENP-A^Nuc^ reconstituted with 601 Widom DNA (147 bp). While no binding was observed between CCAN^ΔT^ and H3^Nuc^, CCAN^ΔT^ bound to CENP-A^Nuc^ only in the presence of CENP-C^N^. (**B**) Comparative SEC profiles of CCAN binding to either H3^Nuc^ or CENP-A^Nuc^ reconstituted with 177-601 Widom DNA (177 bp) and their corresponding Coomassie-blue-stained SDS-PAGE gels. CCAN^ΔT^ bound equally well to both H3^Nuc^ and CENP-A^Nuc^ but only binding to CENP-A^Nuc^ was enhanced in the presence of CENP-C^N^. (**C**) EMSA assays to assess nucleosome binding affinity between either CCAN^ΔCT^ or CCAN^ΔT^ and either H3^Nuc^ or CENP-A^Nuc^ reconstituted with 177-601 Widom DNA (177 bp). The nucleosome was visualized using ethidium bromide. The fraction of bound nucleosome was quantified across three independent replicates as described in the Methods and plotted as mean value with one standard deviation. (**D**) Reconstitution of the CCAN-CENP-A^Nuc^ complex reconstituted with AS1 DNA sequence (αSat171 based) as assessed by SEC and Coomassie-blue-stained SDS-PAGE gels. CCAN^ΔC^ readily bound to CENP-A^Nuc^, and further shift is observed upon addition of CENP-C^N^. (**E**) CCAN^ΔC^ did not detectably bind to CENP-A^Nuc^ reconstituted with αSat147 DNA sequence. The binding was restored in the presence of CENP-C^N^. All reconstitutions were performed in at least two independent duplicates.

**Fig. S11.**
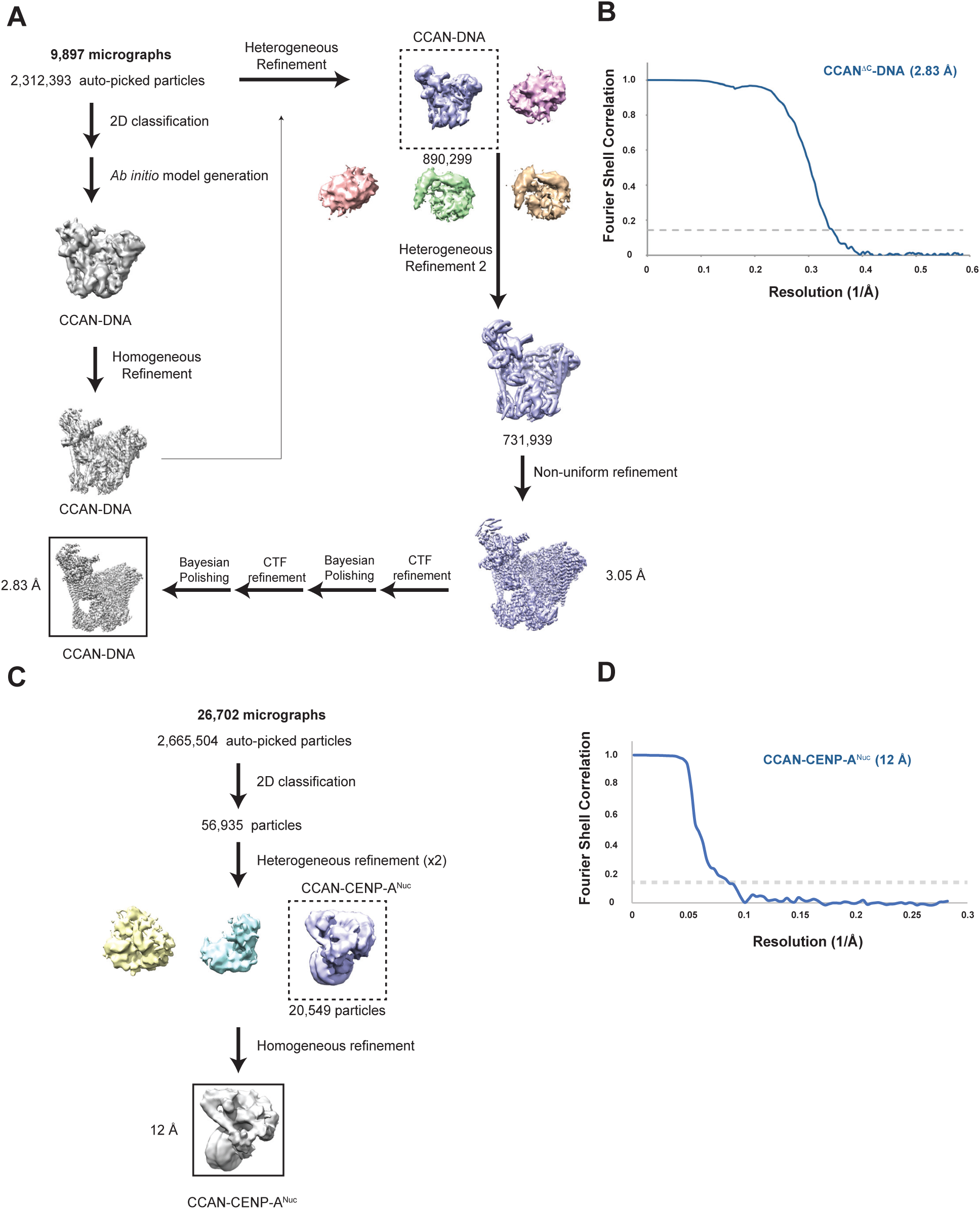
Workflow for cryo-EM reconstruction of CCAN^ΔC^-DNA. (**A**) Work-flow for reconstruction of CCAN^ΔC^-DNA for which 9,897 micrographs were used. (**B**) FSC curve for the CCAN^ΔC^-DNA complex. (**C**) Cryo-EM data processing workflow for the CCAN-CENP-A^Nuc^ AS2 complex (**D**) FSC curve for the CCAN-CENP-A^Nuc^ AS2 complex.

**Table S1.** Cryo-EM data collection, refinement and validation statistics.

|  | apoCCAN<sup>ΔCT</sup> | CENP-A<sup>AS1-Nuc</sup><br>CENP-C<sup>N</sup> | CCAN<sup>ΔT</sup><br>DNA | CCAN<sup>ΔT</sup><br>CENP-A<sup>Nuc</sup> | CCAN<sup>ΔC</sup><br>DNA | CENP-A<sup>AS2-Nuc</sup><br>CENP-C<sup>N</sup> | CCAN<br>CENP-A<sup>Nuc</sup> |
| --- | --- | --- | --- | --- | --- | --- | --- |
| **Data collection and processing** |  |  |  | Dataset1,<br>Dataset2 |  |  |  |
| Magnification | 105,000 | 105,000 | 105,000 | 105,000 85,000 | 105,000 | 105,000 | 105,000 |
| Voltage (kV) | 300 | 300 | 300 | 300 300 | 300 | 300 | 300 |
| Electron exposure (e-/Å<sup>2</sup>) | 50 | 50 | 50 | 50 40 | 40 | 40 | 40 |
| Defocus range (μm) | 1.5-3.0 | 1.5-3.0 | 1.5-3.0 | 1.5-3.0 1.2-2.6 | 1.5-3 | 1.5-3 | 1.2-2.6 |
| Pixel size (Å) | 0.86 | 0.831 | 0.831 | 0.831 0.93 | 0.853 | 0.853 | 0.853 |
| Symmetry imposed | C1 | C1 | C1 | C1 C1 | C1 | C1 | C1 |
| Initial particle images (no.) | 3,797,577 | 2,974,740 | 2,974,740 | 6,876,781<br>(combined) | 2,312,393 | 2,540,375 | 2,665,504 |
| Final particle images (no.) | 569,755 | 1,093,274 | 123,677 | 52,144<br>(combined) | 731,939 | 1,892,204 | 20,549 |
| Map resolution (Å) | 3.2 | 2.7 | 4.5 | 8.9 | 2.83 | 2.44 | 12 |
| FSC threshold | 0.143 | 0.143 | 0.143 | 0.143 | 0.143 | 0.143 | 0.143 |
| Map resolution range (Å) | 3.2-20 | 2.7-20 | 4.5-20 | 7.3-20 | 2.83-20 | 2.44-20 | 8-20 |
| **Refinement** |  |  |  |  |  |  |  |
| Initial model used (PDB code) | *Ab initio* | 6MUP | *Ab initio* | 6MUP, *ab initio* | *Ab initio* | 6MUP | *Ab initio* |
| Model resolution (Å) | 3.2 | 2.7 | 4.5 | 8.9 | 2.83 | 2.44 | 12 |
| FSC threshold |  |  |  |  |  |  |  |
| Model resolution range (Å) | 3.2-20 | 2.7-20 | 4.5-20 | 8.9-20 | 2.83-20 | 2.44-20 | 12-10 |
| Map sharpening *B* factor (Å<sup>2</sup>) | -50 | - 76.6 | - 222.4 | -20 | -30 | -50 | -100 |
| Model composition |  |  |  |  |  |  |  |
| Non-hydrogen atoms | 17406 | 11192 | 18771 | 29822 | 26873 | 11627 | 38871 |
| Protein residues | 2184 | 779 | 2184 | 2963 | 3126 | 779 | 3940 |
| Nucleotide | 0 | 245 | 67 | 305 | 82 | 266 | 352 |
| *B* factors (Å<sup>2</sup>) |  |  |  |  |  |  |  |
| Protein | 125.85 | 185.34 | 173.05 | 182.65 | 83.32 | 185.39 | 118.86 |
| Nucleotide |  | 205.96 | 507.76 | 268.55 | 245.18 | 187.14 | 171.24 |
| R.m.s. deviations |  |  |  |  |  |  |  |
| Bond lengths (Å) | 0.002 | 0.010 | 0.004 | 0.007 | 0.002 | 0.003 | 0.009 |
| Bond angles (°) | 0.494 | 0.922 | 0.939 | 0.933 | 0.465 | 0.524 | 1.025 |
| Validation |  |  |  |  |  |  |  |
| MolProbity score | 1.30 | 1.72 | 1.78 | 1.75 | 1.45 | 1.39 | 1.63 |
| Clashscore | 5.47 | 8.23 | 13.48 | 11.29 | 5.94 | 5.41 | 9.62 |
| Poor rotamers (%) | 0.00 | 0.31 | 0.21 | 0.23 | 0.82 |  | 0.32 |
| Ramachandran plot |  |  |  |  |  |  |  |
| Favored (%) | 97.97 | 96.05 | 97.27 | 96.91 | 97.32 | 97.5 | 97.35 |
| Allowed (%) | 2.03 | 3.95 | 2.73 | 3.04 | 2.61 | 2.5 | 2.46 |
| Disallowed (%) | 0.00 | 0.00 | 0.00 | 0.03 | 0.07 | 0.00 | 0.18 |

**Table S2.**
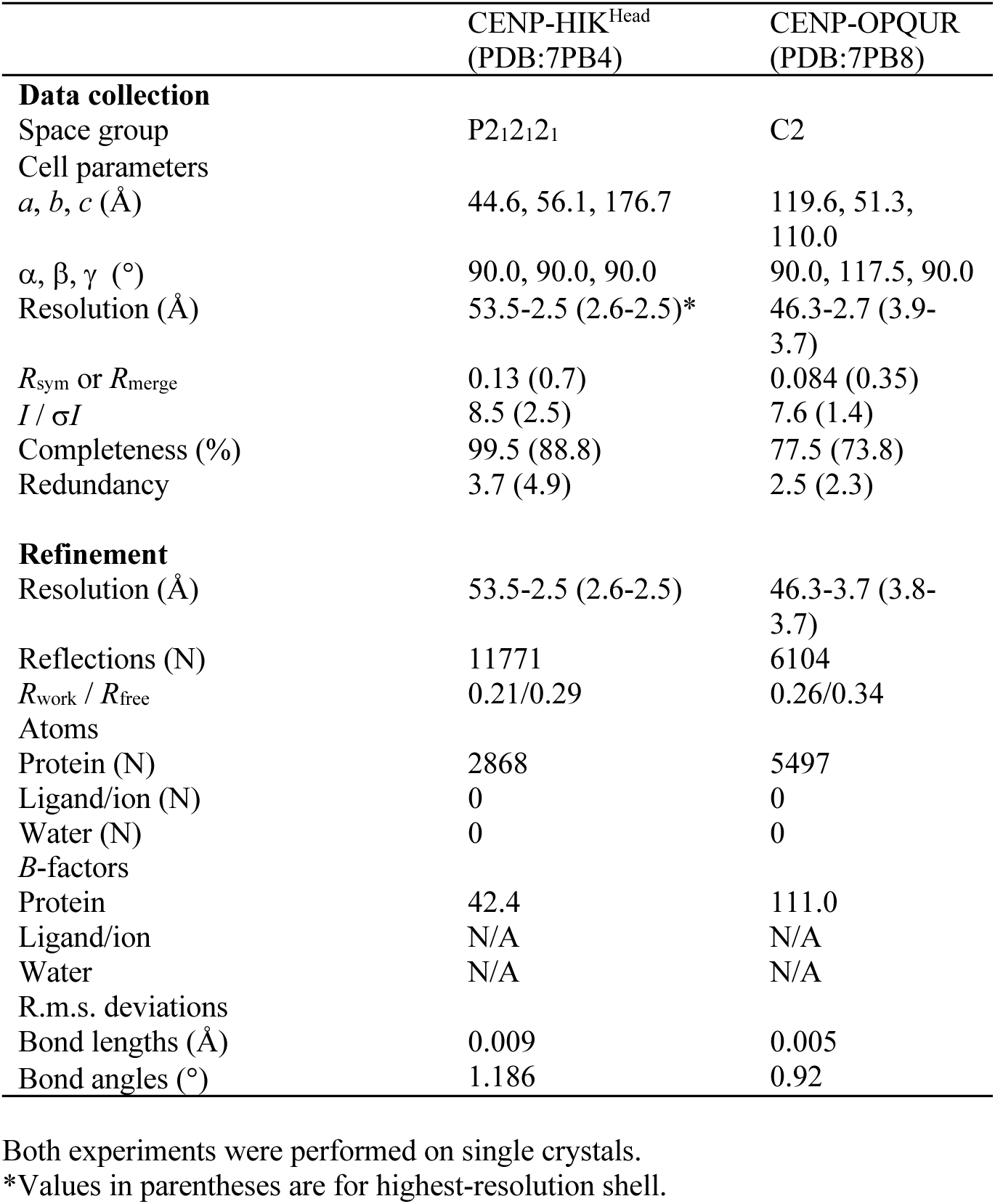
Data collection and refinement statistics (molecular replacement)

|  | CENP-HIK<sup>Head</sup><br>(PDB:7PB4) | CENP-OPQUR<br>(PDB:7PB8) |
| --- | --- | --- |
| **Data collection** |  |  |
| Space group | P2<sub>1</sub>2<sub>1</sub>2<sub>1</sub> | C2 |
| Cell parameters<br>*a*, *b*, *c* (Å) | 44.6, 56.1, 176.7 | 119.6, 51.3,<br>110.0 |
| $\alpha$, $\beta$, $\gamma$ (°) | 90.0, 90.0, 90.0 | 90.0, 117.5, 90.0 |
| Resolution (Å) | 53.5-2.5 (2.6-2.5)* | 46.3-2.7 (3.9-3.7) |
| *R*<sub>sym</sub> or *R*<sub>merge</sub> | 0.13 (0.7) | 0.084 (0.35) |
| *I* / $\sigma I$ | 8.5 (2.5) | 7.6 (1.4) |
| Completeness (%) | 99.5 (88.8) | 77.5 (73.8) |
| Redundancy | 3.7 (4.9) | 2.5 (2.3) |
| **Refinement** |  |  |
| Resolution (Å) | 53.5-2.5 (2.6-2.5) | 46.3-3.7 (3.8-3.7) |
| Reflections (N) | 11771 | 6104 |
| *R*<sub>work</sub> / *R*<sub>free</sub> | 0.21/0.29 | 0.26/0.34 |
| Atoms |  |  |
| Protein (N) | 2868 | 5497 |
| Ligand/ion (N) | 0 | 0 |
| Water (N) | 0 | 0 |
| *B*-factors |  |  |
| Protein | 42.4 | 111.0 |
| Ligand/ion | N/A | N/A |
| Water | N/A | N/A |
| R.m.s. deviations |  |  |
| Bond lengths (Å) | 0.009 | 0.005 |
| Bond angles (°) | 1.186 | 0.92 |

Both experiments were performed on single crystals.
\*Values in parentheses are for highest-resolution shell.

**Movie S1.**

3D variability analysis demonstrates a continuous conformational heterogeneity of the DNA termini of the CENP-A^Nuc^.

**Movie S2.**

Overview of the cryo-EM reconstruction and molecular model of the CCAN^ΔC^-DNA complex.

## Notes

### Competing Interest Statement

The authors have declared no competing interest.

